# Isoprenylcysteine carboxylmethyltransferase-based therapy for Hutchinson–Gilford progeria syndrome

**DOI:** 10.1101/2020.07.23.217257

**Authors:** Beatriz Marcos-Ramiro, Ana Gil-Ordóñez, Nagore I. Marín-Ramos, Francisco J. Ortega-Nogales, Moisés Balabasquer, Pilar Gonzalo, Loïc Rolas, Anna Barkaway, Sussan Nourshargh, Vicente Andrés, Mar Martín-Fontecha, María L. López-Rodríguez, Silvia Ortega-Gutiérrez

## Abstract

Progerin is a mutant prelamin A variant that causes Hutchinson–Gilford progeria syndrome (HGPS, progeria), a rare genetic disease characterized by premature aging and death in childhood. Although several therapeutic approaches have been explored in experimental models, clinical trials have shown very limited benefits in HGPS patients. Here, we describe the development of UCM-13207, a new potent inhibitor of isoprenylcysteine carboxylmethyltransferase (ICMT) that reduces progerin nuclear accumulation and ameliorates the typical alterations in progeroid human and mouse cells. UCM-13207 also improves phenotypic anomalies and extends lifespan in progerin-expressing *Lmna*^*G609G/G609G*^ mice. These results support the potential use of UCM-13207 as a new treatment for progeria.

---

Hutchinson–Gilford progeria syndrome (HGPS), or progeria, is an extremely rare disorder in children, characterized by premature aging and death at an average age of 14.6 years.^1–3^ The molecular cause of HGPS is an autosomal spontaneous dominant point mutation in the *LMNA* gene (encoding nuclear A-type lamins)^4,5^ that leads to the synthesis of progerin, a permanently farnesylated and methylated truncated version of prelamin A.^6^ This mutant protein accumulates abnormally in the inner nuclear membrane causing the multiple and characteristic cellular and organismal alterations of the disease.^1,2^ Evidence exists that reducing progerin accumulation could improve HGPS progression.^7^ In this context, we envisioned that inhibition of isoprenylcysteine carboxylmethyltransferase (ICMT), the enzyme responsible for progerin carboxymethylation, could impact progerin’s affinity by the nuclear membrane and induce its delocalization. It has been reported that genetic blockade of ICMT ameliorates disease phenotype in the Zmpste24-null mouse model of progeria and in HGPS human cells.^8^ In this work we report the development of a new potent and selective ICMT inhibitor (compound **21**, UCM-13207), which significantly improved the hallmarks of progeria. In cells from progeroid mice, treatment with UCM-13207 increased cell viability, delocalized progerin from the nuclear membrane while simultaneously diminished its total levels, and decreased DNA damage. In vivo, it increased body weight, enhanced grip strength, extended lifespan by 20% and induced a decrease in progerin levels in cardiac tissue. Moreover, in human HGPS fibroblasts, compound UCM-13207 increased proliferation rate, induced delocalization of progerin, stimulated proliferative pathways, and delayed overall senescence. Collectively, these findings indicate that inhibition of ICMT could be an effective translational therapy for progeria.

HGPS is a fatal disease that does not yet have a single drug approved for its treatment. To date, lonafarnib, an inhibitor of the enzyme farnesyltransferase, aimed at decreasing progerin farnesylation, is the only compound that has reached clinical trials. However, the outcome of lonafarnib clinical trial has shown only very limited benefit,^9^ probably due to the possibility of alternative prenylation of prelamin A and progerin by geranylgeranyl transferase upon farnesyltransferase inhibition;^10^ however, combination therapies inhibiting both enzymes showed little efficacy when applied to HGPS children.^11^ Other alternative approaches rely on the use of the CRISP/Cas9-based therapy^12,13^ or in microbiome-based interventions,^14^ but these therapies cannot be easily translated to humans yet. Finally, a moderate extension of longevity (by 12%) has been reported in the progerin-expressing *Lmna*^*G609G/G609G*^ mouse model after increasing the levels of ATP and pyrophosphate, but the relevance of this finding to human HGPS remains to be addressed.^15^ Hence, there is clear and urgent need for better therapeutic strategies. In this context, we envisioned that ICMT inhibition, an approach previously not explored in this setting, could avoid the carboxymethylation of progerin and consequently, induce its delocalization from the nuclear membrane.

In our research group, we had previously carried out an extensive medicinal chemistry program aimed at the development of new ICMT inhibitors.^16,17^ From this previous research, we selected a new compound (**1**) as our initial hit, considering that it was able to effectively inhibit ICMT (80% inhibition at 50 μM) and showed low cellular toxicity (cellular viability > 80% at 10 μM). Our findings suggested that the 3-amino-*N*-phenylpropanamide moiety as well as the *n*-octyl chain of compound **1** were key elements for interaction with ICMT.^17^ Accordingly, we envisioned that the phenyl group could be further explored for optimizing cellular viability without significantly compromising ICMT inhibition (Fig. 1a). This focused library was synthesized as detailed in the Supplementary Information. In all cases, the obtained structural data were in agreement with the proposed structures (see Supplementary Information for details). All new compounds were screened for ICMT inhibition. In addition, those compounds which blocked >60% of ICMT activity were assayed for cellular viability in mouse wild-type fibroblasts and in progerin-expressing fibroblasts (*Lmna*^*G609G/G609G*^) (Supplementary Table 1). All compounds that showed the strongest capacity to stimulate the viability of progeroid fibroblasts (values >70% at a 2 μM) were selected for further biological assays with the exception of the furan-containing derivative **13**, because of the narrow efficacy window observed for this compound when tested at higher concentration (Extended Data Fig. 1). Hence, we sought to confirm whether compounds **14**, **17**, and **21** could counteract some of the most characteristic molecular defects found in both progeroid mouse and human cells, including growth arrest, progerin accumulation in the perinuclear rim, Akt inhibition, and increased level of phosphorylated histone 2AX (pH2AX). We found that the three compounds augmented the proliferation rate of progeroid mouse and human cells, with compound **21** yielding the best results in progeroid mouse cells (Fig. 1b) and compound **14** in progeroid human cells (Fig. 1c). Consistent with their ICMT inhibitory activity, all three compounds induced in human HGPS cells a significant delocalization of progerin from the nuclear rim (Fig. 1d, left plot), consistent with their ICMT inhibitory activity. This effect was accompanied by a decrease in the total levels of progerin, as shown in both immunofluorescence experiments in human HGPS cells (Fig. 1d, right plot), and in western blot analysis in mouse progeroid cells (Fig. 1e). A similar trend was noted in the context of cellular proliferation in that although the three compounds induced comparable effects, derivative **14** remained slightly superior in human cells whilst the strongest effects in mouse cells were exerted by compound **21**. Furthermore, and consistent with the increase in cellular proliferation, phospho-Akt levels were higher in treated progeroid cells (Fig. 1f). In addition, the phosphorylated levels of histone H2AX, a marker of nuclear damage associated with aging,^18^ were also significantly reduced in the presence of compounds **14**, **17**, and **21**. Finally, no significant effect was observed on the number of misshapen nuclei in cells treated with this compounds (Extended Data Fig. 2). This is in agreement with the results reported in the ICMT genetically deficient mouse model.^8^ Taken together, these results demonstrate that the new ICMT inhibitors described herein significantly ameliorate the main cellular hallmarks of HGPS in both progeroid fibroblasts from both *Lmna*^*G609G/G609G*^ mice and HGPS patients, suggesting their potential for the treatment of the disease.

**Fig. 1.**
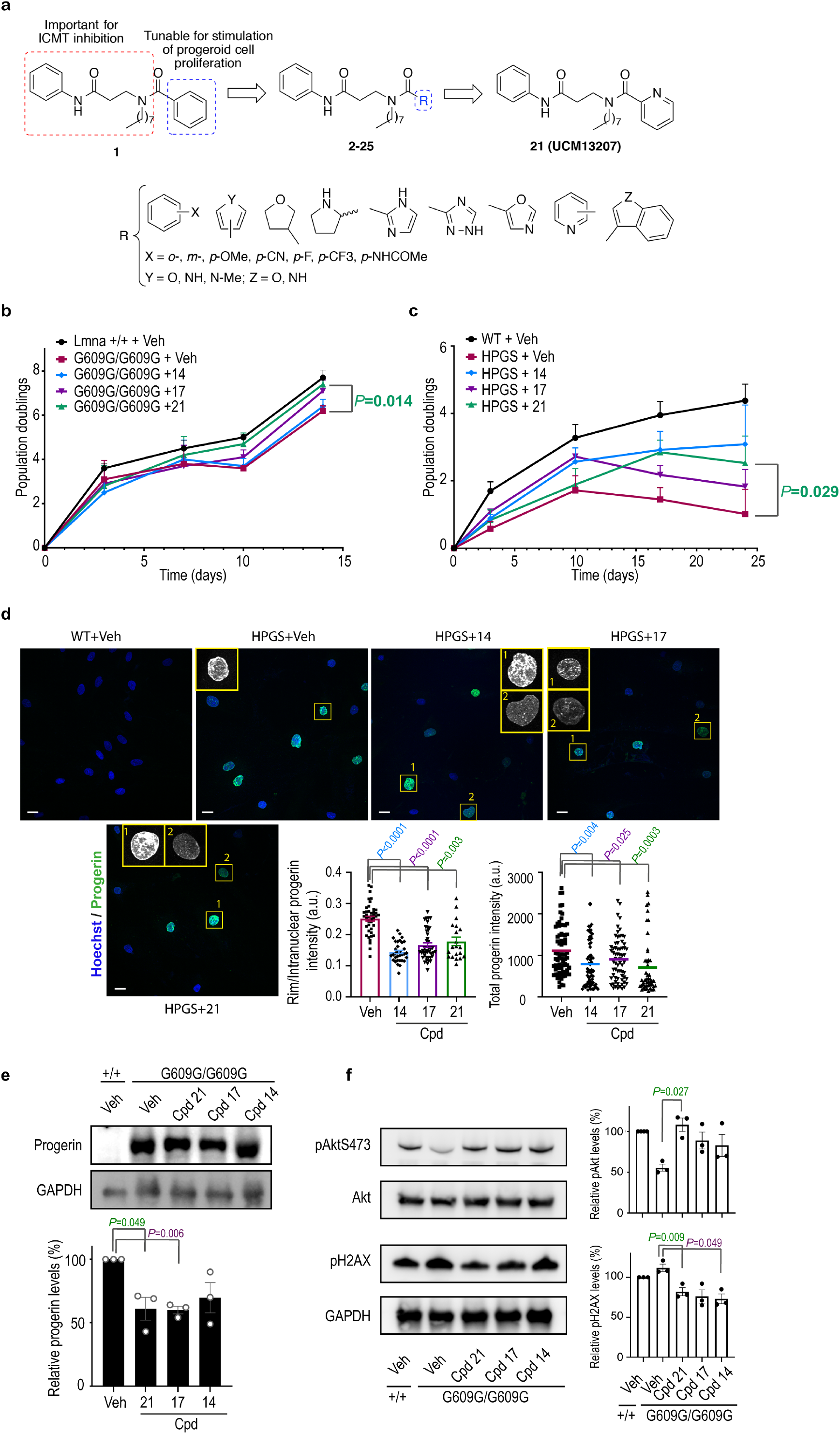
New ICMT inhibitors improve progeria hallmarks in mouse and human HGPS cells. **a**, Design of new ICMT inhibitors. **b**, Growth curves of *Lmna*^*+/+*^ and *Lmna*^*G609G/G609G*^ mice fibroblasts incubated with 0.1% DMSO (vehicle, veh) or ICMT inhibitors at 10 μM during 14 days (n≥4 independent experiments; two-tailed Student’s t-test). **c**, Growth curves of human wild type (WT) or HGPS fibroblasts incubated with vehicle or different ICMT inhibitors at 2 μM during 24 days (n≥4 independent experiments; two-tailed Student’s t-test). **d**, Immunofluorescence images of human WT or HPGS fibroblasts treated with vehicle or different ICMT inhibitors at 2 μM during 17 days and stained with anti-progerin (green) and Hoechst (blue). Magnification shows subnuclear distribution of progerin, that was delocalized from the nuclear rim towards the nucleoplasm (1, quantification in middle plot) and/or decreased (2, quantification in right plot) after treatment with ICMT inhibitors (at least 47 cells quantified per condition from ≥2 independent experiments; two-tailed Student’s t-test). Scale bar: 20 μm. **e**, Immunoblot of progerin from *Lmna*^*+/+*^ or *Lmna*^*G609G/G609G*^ mouse fibroblasts incubated with vehicle or ICMT inhibitors during 14 days. Protein levels were normalized against total GAPDH (n≥3 independent experiments; plot shows mean±sem, two-tailed Student’s t-test). **f**, Immunoblot and quantification of phosphorylated Akt (pAkt) and phosphorylated histone 2AX (pH2AX) from *Lmna*^*+/+*^ or *Lmna*^*G609G/G609G*^ mouse fibroblasts incubated with vehicle or ICMT inhibitors during 14 days. Protein levels were normalized against total Akt or GAPDH respectively (n≥3 independent experiments; plot shows mean±sem, two-tailed Student’s t-test).

**Fig. 2.**
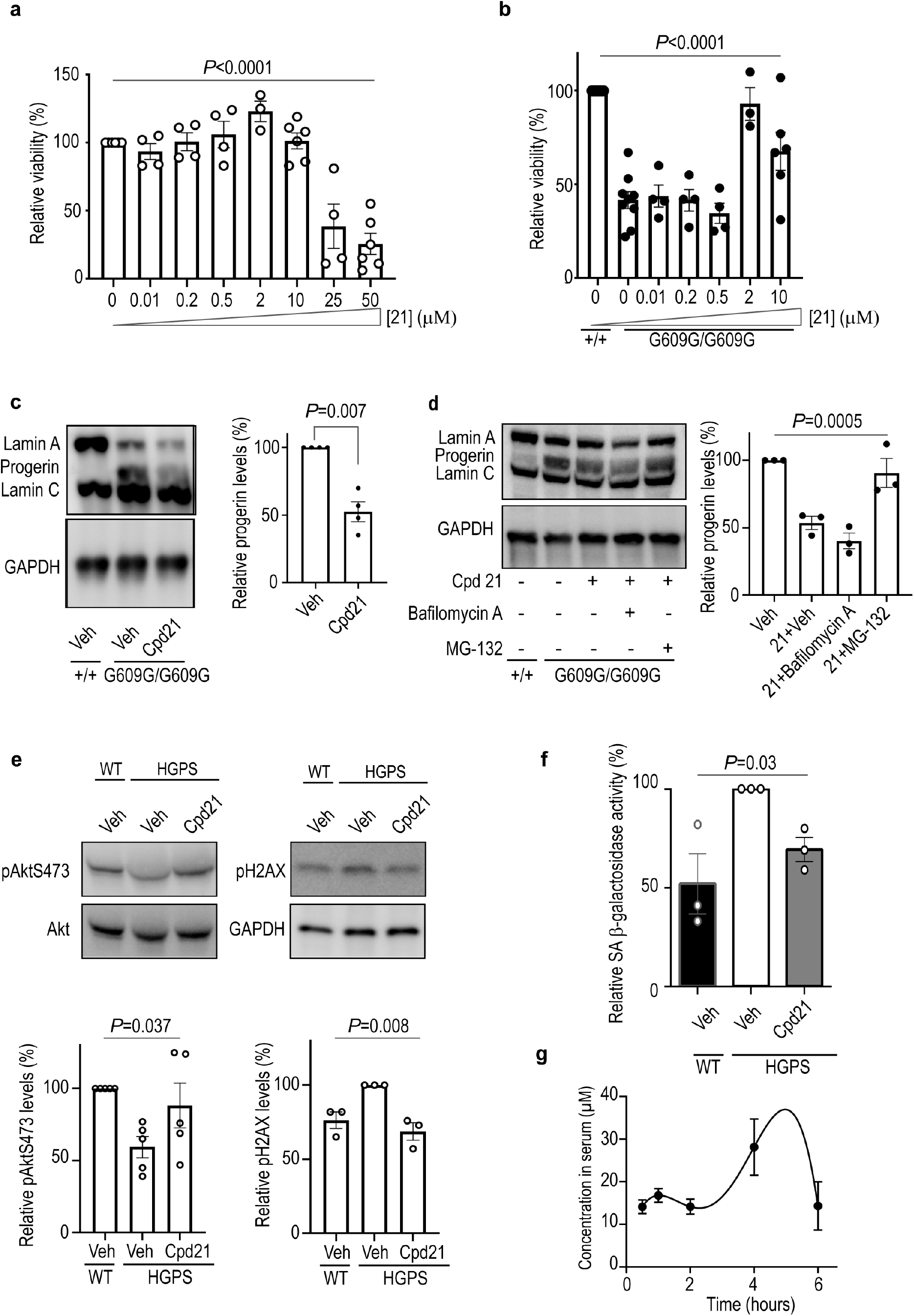
Compound 21 ameliorates progeria phenotype in mice and human HGPS. **a**, *Lmna*^*+/+*^ fibroblasts were incubated with compound **21** at different concentrations during 72 h and viability was determined by an MTT assay (n≥4 independent experiments; one-way ANOVA). **b**, *Lmna*^*+/+*^ or *Lmna*^*G609G/G609G*^ fibroblasts were incubated with compound **21** during 72 h at different concentrations and viability was assessed by MTT assay (n≥4 independent experiments; one-way ANOVA). **c**, Cropped immunoblot of lamin A, progerin, and lamin C from *Lmna*^*+/+*^ or *Lmna*^*G609G/G609G*^ fibroblasts incubated with 0.1% DMSO (vehicle, veh) or compound **21** at 2 μM during 14 days (n≥3 independent experiments; plot shows progerin level±sem, two-tailed Student’s t-test). **d**, Cropped immunoblot and quantification of lamin A, progerin and lamin C from *Lmna*^*+/+*^ or *Lmna*^*G609G/G609G*^ fibroblasts. Cells were incubated with vehicle or compound **21** at 2 μM during 14 days and then bafilomycin A (24 h at 25 nM) or MG-132 (5 h at 5 μM) was added when indicated (n=3 independent experiments; plot shows mean±sem, one-way ANOVA). **e**, Cropped immunoblot and quantification of phosphorylated Akt (pAkt) and phosphorylated histone 2AX (pH2AX) from wild type (WT) or HPGS human fibroblasts incubated with vehicle or compound **21** at 2 μM during 10 days. Protein levels were normalized against total Akt or GAPDH (n≥3 independent experiments; plot shows mean±sem, one-way ANOVA). **f**, Human WT or HGPS fibroblasts were incubated with vehicle or compound **21** at 2 μM during 15 days. The activity of senescence-associated (SA) β-galactosidase was determined by incubating the cells with fluorescein di-β-D-galactopyranoside (FDG) for 24 h and measuring the resultant fluorescein production (n≥3 independent experiments in triplicates; plot shows mean±sem, two-tailed Student’s t-test). **g**, WT C57BL/6 mice (n=2 females; n=2 males per time point) were treated with compound **21** (intraperitoneal injection, 40 mg/kg), blood was extracted at different time points and compound concentration in serum determined by high-performance liquid chromatography coupled to mass spectrometry.

Among the three tested compounds, derivative **21 (UCM-13207)** systematically stood out as the one inducing the strongest effects in all the performed experiments. Hence, it was selected for further exploration regarding its in vitro toxicity-efficacy window and the main pharmacokinetic parameters before assessing its in vivo efficacy in a mouse model of progeria. UCM-13207 did not cause appreciable cellular toxicity at least up to 10 μM (Fig. 2a), whereas its efficacy to preserve the proliferation of progeroid cells was dose-dependent, with 2 μM showing the maximal effect (Fig. 2b). Consistent with these results, compound **21** at 2 and 10 μM induced a decrease in progerin level (Fig. 2c and 1e, respectively) without affecting in a significant manner the levels of wild-type lamin A and lamin C (Extended Data Fig. 3). The decrease in progerin levels is mediated by the proteasome pathway since blocking this complex with the specific inhibitor MG-132 reversed the effects of compound **21**, whereas treatment with the lysosome pathway inhibitor bafilomycin A did not significantly affect progerin downregulation induced by compound **21** (Fig. 2d). Moreover, human HGPS fibroblasts treated with 2 μM compound **21** significantly increased phospho-Akt and reduced phospho-H2AX levels (Fig. 2e) and diminished the levels of senescence-associated (SA) β-galactosidase activity (Fig. 2f), a biomarker of senescent cells.^19^

**Fig. 3.**
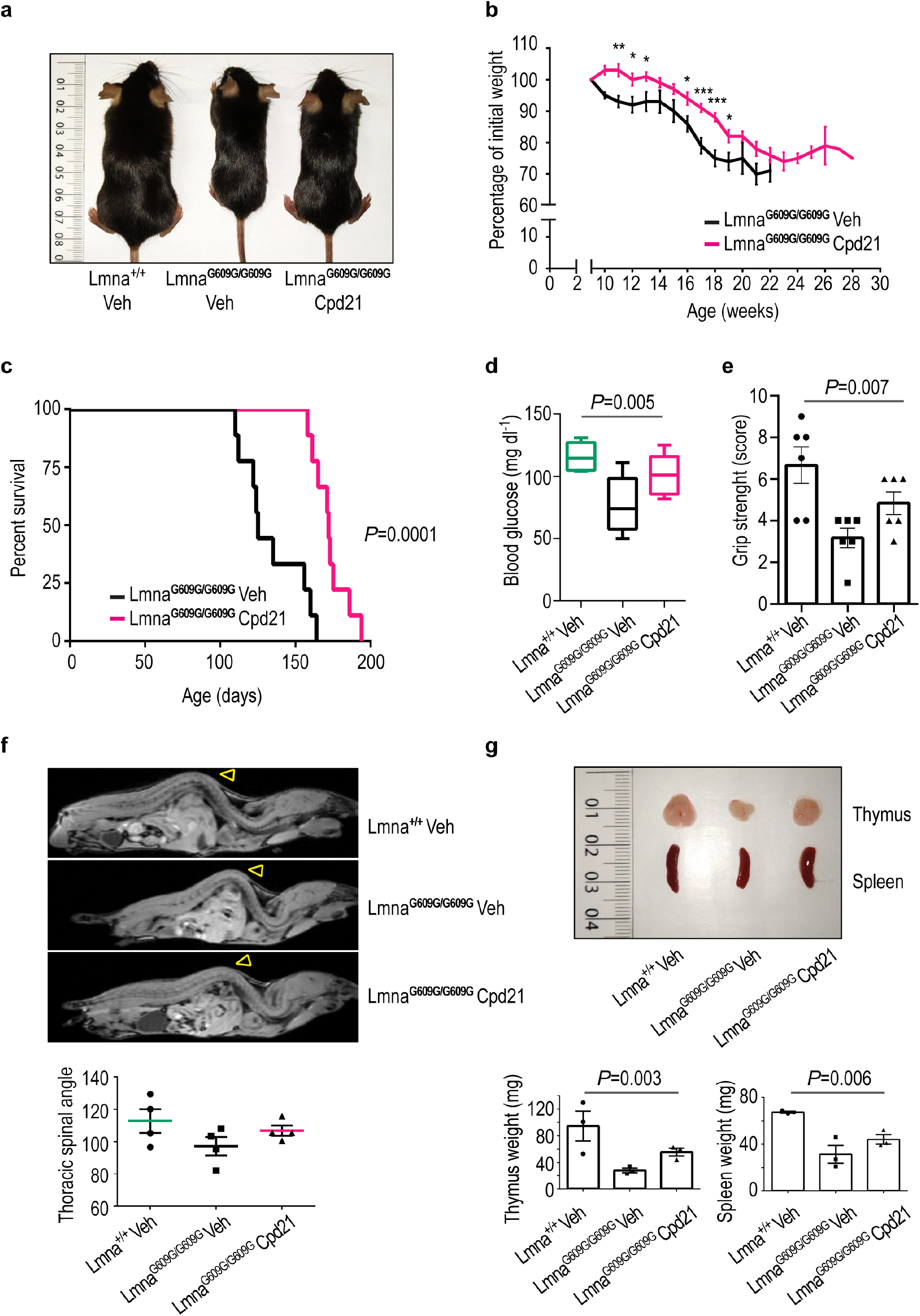
Improvement of progeroid phenotype in vivo upon treatment with compound 21. **a**, Representative photograph of a 3-month-old *Lmna*^*+/+*^ mouse, a *Lmna*^*G609G/G609G*^ mouse and a *Lmna*^*G609G/G609G*^ mouse treated with compound **21** (40 mg/kg). **b**, Body weight versus age plot of *Lmna*^*G609G/G609G*^ mice treated with vehicle (n=6) or with compound **21** (n=9) (plot shows mean±sem, two-tailed Student’s t-test, p-value *<0.05, **<0.005, ***<0.0005). **c**, Kaplan-Meier survival plot showing a significant increase in life span in *Lmna*^*G609G/G609G*^ mice treated with compound **21** (n=9) compared with *Lmna*^*G609G/G609G*^ mice treated with vehicle (n=9) (P=0.0001, log-rank/Mantel-Cox test). **d**, Glycemia in *Lmna*^*+/+*^ and *Lmna*^*G609G/G609G*^ mice treated with compound **21** or vehicle (n=6 per condition). Data are represented by box plots and whiskers are minimum to maximum values (one-way ANOVA). **e**, Grip strenght in *Lmna*^*+/+*^ and *Lmna*^*G609G/G609G*^ mice treated with compound **21** or vehicle (n=5 per condition, one-way ANOVA). **f**, Resonance of a 3-month-old *Lmna*^*+/+*^ mouse, a vehicle-treated *Lmna*^*G609G/G609G*^ mouse and a compound **21**-treated *Lmna*^*G609G/G609G*^ mouse. Note that the marked curvature of the spine called lordokyphosis that is characteristic of progeria is reduced upon treatment with compound **21**. Plot represents the average of inner column angle (yellow arrowhead) of ≥ 4 mice per condition. **g**, Representative photographs of thymus and spleen from a 3-month-old *Lmna*^*+/+*^ mouse, a vehicle-treated *Lmna*^*G609G/G609G*^ mouse and a compound **21**-treated *Lmna*^*G609G/G609G*^ mouse. Plots represent average of thymus or spleen weight from ≥ 3 mice per condition.

With respect to the pharmacokinetic parameters, we first determined the cell culture and serum stability, the intrinsic clearance, and the serum free drug or fraction unbound (Fu). Our in vitro stability results indicated excellent values for the compound half-life in both human and mouse cell culture media as well as in human serum (t_1/2_>24 h) although a more moderate value was obtained in the case of mouse serum (t_1/2_=29±2 min). The in vitro microsomal intrinsic clearance was 40±6 μL/min/μg protein in human samples and 144±29 μL/min/μg protein in mouse samples. These values are predictive of a medium and high in vivo clearance, respectively, but the fact that the Fu of compound **21** was 0.02 (with a *K*_D_ value for human serum albumin of 7.2 μM) might reduce the metabolic clearance by the liver and, in turn, increase its half-life. Indeed, the in vivo half-life curve after intraperitoneal (ip) administration showed a significant and sustained concentration of the compound up to 6 h post-administration, reaching a maximum concentration close to 40 μM after 5 h (Fig. 2g). In addition, neither signs of acute toxicity nor tissue or pathological damage was observed at doses up to 80 mg/kg of compound **21**.

Having demonstrated the in vitro efficacy of compound **21** in mouse and human progeroid cells, we evaluated its therapeutic potential in the *Lmna*^*G609G/G609G*^ progeria mouse model, as it recapitulates the majority of the alterations observed in HGPS patients.^20–23^ Remarkably, progeroid mice treated with the ICMT inhibitor **21** showed significantly improved body weight at all ages tested (Fig. 3a,b) and exhibited increased lifespan (Fig. 3c). The beneficial effects of compound **21** were observed in both males and females (Extended Data Fig. 4–6). Moreover, no signs of toxicity were observed in wild-type mice receiving the same dose of compound (Extended Data Fig. 7). Encouraged by these results, we explored other phenotypic characteristics in *Lmna*^*G609G/G609G*^ mice, including reduced serum glucose levels, muscular weakness, bone defects, and marked involutions of thymus and spleen.^20^ Treatment of progeroid mice with compound **21** increased serum glucose to levels close to those in wild-type controls (Fig. 3d), remarkably improved both grip strength and lordokyphosis (abnormal convexity in the curvature of the spine when viewed from the side) (Fig. 3e,f), and significantly increased the size of thymus and spleen (Fig. 3g) when compared to untreated mice. In addition, recent evidence points to progerin expression in cardiac tissue as the main cause of the cardiovascular alterations that ultimately lead to death in progeria patients.^22,24^ Hence, we sought to confirm whether treatment with compound **21** was able to reduce progerin levels in vivo, and we found that this is indeed the case, as treated *Lmna*^*G609G/G609G*^ mice showed decreased levels of progerin in the thoracic aorta, in the aortic arch, and in myocardial tissue (Extended Data Fig. 8).

In conclusion, we report herein a novel strategy to reduce the anomalous accumulation of progerin in the nuclear rim membrane, which is considered the main cause of the fatal phenotype developed by HGPS patients. Specifically, we show that a new ICMT inhibitor, compound **21** or UCM-13207, significantly induces the mislocalization of progerin and reduces its levels, leading to a substantial overall amelioration of the cellular hallmarks of progeria in both mouse and human cells.

Importantly, treatment with compound **21** significantly improved the phenotype of progeric *Lmna*^*G609G/G609G*^ mice, including lifespan extension in both males and females. Collectively, these findings have important clinical implications in that they validate the clinical relevance of ICMT as a promising therapeutic target for progeria treatment, providing a new pharmacological strategy in a field dramatically devoid of validated therapeutic targets beyond farnesyltransferase. Furthermore, our results pave the way for targeting ICMT as a new pharmacological strategy for succeeding in the long sought objective of making a meaningful difference to the lives of HGPS patients.

## Online content

Any methods, additional references, Nature Research reporting summaries, source data, and statements of data availability are available upon request.

## Acknowledgements

This work was supported by grants from The Progeria Research Foundation (PRF 2016-65) and the Spanish MINECO (SAF2016-78792-R, PID2019-106279RB-I00). Authors thank Fundación La Caixa, CEI Moncloa, and MINECO for predoctoral fellowships to A.G., N.I.M.-R., and F.J.O.-N. and M.B., respectively. The authors thank C. López-Otín for kindly donating *Lmna*^*G609G/G609G*^ progeroid and their corresponding wild-type fibroblasts, and to UCM’s CAIs Cytometry and Fluorescence Microscopy, Genomics, NMR, and Mass Spectrometry for their assistance. The CNIC is supported by the Ministerio de Ciencia e Innovación, the Instituto de Salud Carlos III, and the pro-CNIC Foundation, and is a Severo Ochoa Center of Excellence (grant SEV-2015-0505). The generation of the anti-progerin antibody was funded by the Wellcome Trust (098291/Z/12/Z to S.N.).

## Author Contributions

B.M., M.M.-F., M.L.L.-R., and S.O.-G. conceived and designed experiments. B.M. performed cellular and in vivo experiments and analyzed data. A.G.-O carried out chemical synthesis, performed pharmacokinetic experiments, and analyzed data. N.I.M.-R. performed initial cellular experiments and analyzed data. F.J.O.-N., and M.B. carried out chemical synthesis. L.R., A.B., and S.N. developed and provided the mouse anti-progerin antibody. V.A. and P.G. provided the progeroid mouse model and performed immunohistochemistry experiments. S.O.-G., M.L.L.-R., M.M.-F., V.A., and B.M. wrote the manuscript. All authors revised the manuscript. The manuscript was written through contributions of all authors. All authors revised the manuscript and have given approval to the final version.

## Additional Information

**Extended data** is available for this paper.

**Supplementary information** is available for this paper.

**Reprints and permissions information** is available for this paper.

**Correspondence and requests for materials** should be addressed to S.O.-G.

## Methods

### Compound synthesis

Unless stated otherwise, starting materials, reagents and solvents were purchased as high-grade commercial products from Sigma-Aldrich, Acros, Abcr, Fluorochem, Scharlab or Honeywell, and were used without further purification. Dichloromethane (DCM) was dried using a Pure Solv™ Micro 100 Liter solvent purification system. All reactions in organic solvents were performed under an argon atmosphere in dry glassware. Analytical thin-layer chromatography (TLC) was run on Merck silica gel plates (Kieselgel 60 F-254) with detection by UV light (254 nm), ninhydrin solution, or 10% phosphomolybdic acid solution in ethanol. Flash chromatography was performed on a Varian 971-FP flash purification system using silica gel cartridges (Varian, particle size 50 μm) or on glass columns using silica gel type 60 (Merck, particle size 230 μm, 400 mesh). All compounds were obtained as oils, except for those whose melting points (mp) are indicated, which were solids. Melting points (mp, uncorrected) were determined on a Stuart Scientific electrothermal apparatus. Infrared (IR) spectra were measured on a Bruker Tensor 27 instrument equipped with a Specac ATR accessory of 5200-650 cm^−1^ transmission range; frequencies (ν) are expressed in cm^−1^. Optical rotation [α] was measured on an Anton Paar MCP 100 modular circular polarimeter using a sodium lamp (λ = 589 nm) with a 1 dm path length; concentrations (c) are given as g/100 mL. NMR spectra were recorded at room temperature on a Bruker DPX 300 (^1^H, 300 MHz; ^13^C, 75 MHz) spectrometer at the Universidad Complutense de Madrid (UCM) NMR facilities. Chemical shifts (δ) are expressed in parts per million relative to internal tetramethylsilane; coupling constants (*J*) are in hertz (Hz). The following abbreviations were used to describe peak patterns when appropriate: s (singlet), d (doublet), t (triplet), q (quartet), qt (quintet), m (multiplet), and br (broad). 2D NMR experiments (HMQC and HMBC) of representative compounds were carried out to assign protons and carbons of the new structures. High resolution mass spectrometry (HRMS) was carried out on a FTMS Bruker APEX Q IV spectrometer in electrospray ionization (ESI) mode at UCM’s spectrometry facilities. Elemental analyses of solid compounds were obtained on a LECO CHNS-932 apparatus at the UCM’s analysis services and were within 0.4% of the theoretical values.

For all final compounds, purity was determined by high-performance liquid chromatography coupled to mass spectrometry (HPLC-MS) using an Agilent 1200LC-MSD VL instrument, and satisfactory chromatograms confirmed a purity of at least 95% for all tested compounds. LC separation was achieved with an Eclipse XDB-C18 column (5 μm, 4.6 mm × 150 mm), together with a guard column (5 μm, 4.6 mm × 12.5 mm) with a flow rate of 0.5 mL/min. The gradient mobile phase consisted of A (95:5 water/acetonitrile) and B (5:95 water/acetonitrile) with 0.1% formic acid as solvent modifier. The gradient started at 0% B (for 5 min), increased linearly to 90% B over the course of 10 min, and up to 100% B for 7 min more, before being kept isocratic at 100% B for 4 min and decreased to 0% B for the final 4 min (total run time = 30 min). Spectra were acquired in positive or negative ionization mode from 100 to 1000 m/z and in UV-mode at four different wavelengths (210, 230, 254, and 280 nm). MS analysis was performed with an ESI source. The capillary voltage was set to 3.0 kV and the fragmentor voltage was set at 72 eV. The drying gas temperature was 350 °C, the drying gas flow was 10 L/min, and the nebulizer pressure was 20 psi. The enantiomeric excess (ee) was determined by HPLC using a chiral column in reversed-phase chromatography. Chiralpak® IA (5 μm, 4.6 mm × 15.0 mm) was used as the stationary phase, equipped with an Eclipse XDC-C18 precolumn (5 μm, 4.6 mm × 12.5 mm) using a flow of 0.5 mL/min. The mobile phase consisted of A (20 mM NH_4_HCO_3_, pH 9) and B (acetonitrile) using an isocratic method of 80% B for 10 (chiral HPLC method A) or 30 min (chiral HPLC method B). HPLC traces were compared to racemic samples obtained by mixing equal amounts of the enantiopure compounds independently obtained. The synthesis and structural characterization of compound **21 (** UCM-13207) is detailed below. Full details regarding the synthetic procedures and characterization data of all compounds are given in the Supplementary Information.

### Synthesis of *N*^2^-octyl-*N*^1^-phenyl-*N*^2^-(pyridin-2-ylcarbonyl)-β-alaninamide (21, UCM-13207)

To a solution of picolinic acid (133 mg, 1.1 mmol) in anhydrous DCM (4 mL/mmol), EDC (207 mg, 1.1 mmol) and HOBt (146 mg, 1.1 mmol) were added. The reaction mixture was stirred at rt for 1 h. Then, a solution of *N*^*2*^-octyl*-N*^*1*^-phenyl-β-alaninamide (150 mg, 0.54 mmol) in anhydrous DCM (2.0 mL/mmol) was added and the reaction mixture was stirred at rt for 16 h. The reaction crude was washed with saturated aqueous solutions of NaHCO_3_ and NaCl, consecutively. The organic extracts were dried over Na_2_SO_4_, filtered, and the solvent removed under reduced pressure. The residue was purified by column chromatography (hexane/EtOAc, 2:8) to yield compound **21** in 92% yield (189 mg).

R_f_ (hexane/EtOAc, 3:7): 0.30. IR (ATR, cm^−1^): 3309 (NH), 1685 (CON), 1615, 1547, 1495, 1443 (Ar). ^1^H-NMR (CDCl_3_, δ): Amide rotamers A:B, 2:1; 0.85 (t, *J* = 6.7 Hz, 3H, CH_3_), 1.11-1.25 (m, 10H, (C*H*_2_)_5_CH_3_), 1.57 (m, 2H, C*H*_2_(CH_2_)_5_CH_3_), 2.81 (t, *J* = 6.4 Hz, 2H, CH_2_CO), 3.37 (t, *J* = 7.4 Hz, 2H, (CH_2_)_6_C*H*_2_N, rotamer A), 3.46-3.47 (m, 2H, (CH_2_)_6_C*H*_2_N, rotamer B), 3.69-3.75 (m, 2H, COCH_2_C*H*_2_N, rotamer B), 3.87 (t, *J* = 6.3 Hz, 2H, COCH_2_C*H*_2_N, rotamer A), 7.05 (t, *J* = 7.1 Hz, 1H, H_4_), 7.24-7.32 (m, 3H, H_3_, H_5_, H_5′_), 7.54-7.56 (m, 3H, H_2_, H_6_, H_4′_), 7.71-7.76 (m, 1H, H_3′_), 8.55 (d, *J* = 4.4 Hz, 1H, H), 8.64 (br s, 1H, NH, rotamer B), 8.92 (br s, 1H, NH, rotamer A). ^13^C-NMR (CDCl_3_, δ): 14.0 (CH_3_), 22.6, 26.5, 28.8, 29.0 (2C), 31.7 ((*C*H_2_)_6_CH_3_), 36.5 (*C*H_2_CO), 43.2 (CH_2_N), 50.0 ((CH_2_)_6_*C*H_2_N), 119.9 (C_2_, C_6_), 123.1, 123.9, 124.4 (C_4_, C_3′_, C_5′_), 128.8 (C_3_, C_5_), 137.0 (C_4′_), 138.4 (C_1_), 148.5 (C_6′_), 154.5 (C_2′_), 169.5, 169.7 (CONH, CON). HRMS (ESI, m/z): Calculated for C H N O Na [M+Na]^+^: 404.2308. Found: 404.2276.

### Determination of ICMT activity

Synthesized compounds were tested for their ability to inhibit human ICMT activity using Sf9 membranes containing the recombinantly expressed enzyme. In this assay, a mixture of biotin-farnesyl-L-cysteine and tritiated S-adenosylmethionine in the presence or absence of the compound under study was prepared and the reaction was initiated by the addition of the Sf9 membrane homogenates. The inhibitory capacity of the compounds is expressed as percentage of inhibition of the methyl esterification step, in which the tritiated methyl group of the methyl donor *S*-adenosylmethionine is transferred to the substrate biotin-farnesyl-L-cysteine as described previously^25^ and the radioactivity incorporated is quantified.

### Cell lines and culture

Progeroid mouse fibroblasts (*Lmna*^*G609/G609*^) and their wild-type counterparts were kindly donated by Prof. Carlos López Otín (Oviedo University, Spain). Cells were grown in Dulbecco’s Modified Eagle medium (DMEM, Invitrogen) supplemented with 10% heat-inactivated fetal bovine serum (FBS, HyClone), 1% L-glutamine (Invitrogen), 1% sodium pyruvate (Invitrogen), 50 U/ mL penicillin and 50 μg/mL streptomycin (Invitrogen). Human progeroid or healthy fibroblasts (HGADFN167, HGADFN003, HGADFN143, and HGFDFN168) were obtained from The Progeria Research Foundation and cultured in 15% DMEM, 50 U/mL penicillin and 50 μg/mL streptomycin. All cells were incubated in a humidified atmosphere at 37 °C in the presence of 5% CO_2_.

### Cell viability and proliferation assays

The effect of the different compounds on the cellular viability was determined through standard MTT assays.^26–28^ Cells were seeded in 96-well plates at a density of 5 × 10^2^ cells per well in the corresponding medium with 10% FBS for 24 h prior to treatments. The medium was then replaced by fresh medium containing different concentrations of compounds tested or by medium containing the equivalent volume of dimethylsulfoxide (DMSO, vehicle control). Cells were treated for 72 h, and then medium was replaced by fresh medium with 2 mg/mL of MTT (3-(4,5-dimethylthiazol-2-yl)-2,5-diphenyltetrazolium bromide, Sigma-Aldrich) and cells were incubated for 4 h at 37 °C in the dark. Once supernatants were removed, formazan crystals previously formed by viable cells were dissolved in DMSO (100 μL/ well) and absorbance was measured at 570 nm (OD570-630) using an Asys UVM 340 (Biochrom Ltd., Cambridge, UK) microplate reader. Background absorbance from blank wells containing only media with compound or vehicle were substracted from each test well.

Measurements were performed in triplicate, and each experiment was repeated at least three times. The viability values were calculated from the percentage of viable cells with respect to 100% viability of the vehicle-treated cells.

For proliferation assays, AF3 mouse fibroblasts (15 × 10^3^ cells) or human progeroid fibroblasts (25 × 10^3^ cells) were seeded on 24-well plates in the corresponding medium. After 24 h, the medium was replaced by fresh medium containing different concentration of compounds or the equivalent volume of DMSO. Every three (mouse cells) or seven (human cells) days, cells were treated with trypsin, counted, and reseeded with fresh medium and compound in increasing plate sizes. Population doublings (PD) were calculated using the formula PD=Log (harvested/seeded)/Log(2).

### Immunoblot analysis

Western blot analysis was carried out as described previously.^29–31^ Mouse or human progeroid cells previously incubated with compound of interest or DMSO were seeded at density of 50 × 10^3^ cells on 24-well plates and incubated for 24 h in the appropriate medium with fresh compound. Then, cells were lysed with cold laemmly buffer (50 mM Tris-HCl pH 7.4, 150 mM NaCl, 1% Igepal) with a mixture of protease and phosphatase inhibitors (1 mM PMSF, 10 mM NaF, 0.3 mg/mL CalyculinA, 1 mM sodium orthovanadate, 10 mM β-glycerol-phosphate; all from Sigma-Aldrich and complete protease inhibitor cocktail, from Roche). Proteins were denatured at 95 °C for 5 min and the lysates were immediately used or stored at −20 °C until their use. Samples were analyzed by electrophoresis in polyacrylamide gels (4-20% SDS-PAGE, BioRad) and the proteins transferred to nitrocellulose membranes (GE Healthcare, Amersham). After 1 h of incubation in blocking buffer [100 mM Tris-HCl pH 8.0, 150 mM NaCl and 0.05% Tween-20 (TBS-T) with 3% of bovine serum albumin (BSA)], blots were incubated overnight at 4 °C with the corresponding primary antibody: anti-phospho-histone 2AX (Ser139) (EMD Millipore, 05-636, 1:1000), anti-phospho-Akt (Cell Signaling, 4060S, 1:1000), anti-Akt (Cell Signaling, 9272, 1:1000), anti-progerin (Nourshargh S. lab, 1:1000), anti-lamin A/C (Santa Cruz Biotechnology, sc-376248, 1:500) and anti-GAPDH (Cell Signaling, 2118S, 1:10000) as antibody for loading control. Then, membranes were washed (3×10 min) with TBS-T and incubated with the corresponding secondary antibody conjugated with peroxidase (Sigma-Aldrich) for 1 h at rt. Membranes were washed (6×5 min) with TBS-T and proteins were visualized by quimioluminiscence (GE Healthcare Amersham™ ECL™ Western Blotting Detection Reagents, Fisher Scientific) in a Fujifilm LAS-3000 imager. The bands were quantified by densitometry using the program ImageJ (NIH).

For progerin degradation studies, mouse progeroid cells previously incubated with compound of interest or DMSO during 14 days were seeded at density of 50 × 10^3^ cells on 24-well plates in the appropriate medium with fresh compound and BrefeldynA (24 h at 25 nM) or MG-132 (5 h at 5 mM) were added. Then, cells were lysed and subjected to immunoblot analysis for progerin as described above.

### Immunocytofluorescence

Human progeroid cells previously incubated with compound of interest or DMSO were seeded at a density of 2 × 10^4^ cells/well over 12 mm diameter slide covers and were incubated for 24 h at 37 °C with compound of interest. Cells were fixed with cold methanol (Sigma-Aldrich) during 15 min and permeabilized with PBS containing 0.1% Triton X-100 (PBS-T, Sigma-Aldrich). Then, cells were blocked with TBS 3% BSA during 20 min at rt, followed by incubation with an anti-progerin antibody (Santa Cruz Biotechnology, SC-81611, 1:500) in TBS with 3% BSA at 37°C for 3 h. After that, cells were washed with TBS (1x) and incubated in absence of light for 1 h with fluorescent goat anti-mouse (1:500, Alexa Fluor 488, Life Technologies) diluted in TBS with 3% BSA. Later, cells were additionally washed with TBS (1x) and incubated with Hoechst 33258 (5 μg/mL, Sigma-Aldrich) for 10 min at rt in order to visualize cellular nucleus. Finally, cells were washed with TBS (2x) and the slide cover was carefully mounted with Immumount (Thermo Scientific). The visualization was carried out using confocal microscopy (Olympus IX83) equipped with a 40X oil immersion lens and the appropriate excitation and emission filters at the microscopy UCM facilities.

Quantification of progerin intensity or internalization was performed using ImageJ (NIH). For quantification of progerin intensity, images taken under the same conditions were processed through the plug-in “split channels”. Then, a threshold was applied to blue Hoescht channel in order to create a region containing each cell nucleus in the image. These regions were extrapolated to progerin channel and green fluorescence intensity was determined for each nucleus. For progerin internalization quantification, the same methodology was followed, but once the nuclei regions were determined for cells positive for progerin, they were made 2 pixels smaller and subtracted from the original region to generate a new region comprising the fluorescence intensity at nucleus periphery. The main pixels intensity of progerin included in this peripheral region was quantified and compared with fluorescence intensity from the whole nuclear area.

### Senescence-associated (SA) β-galactosidase activity

SA β-galactosidase activity was determined by the FDG method.32 Human progeroid cells previously incubated with compound of interest or vehicle were seeded at a density of 5 × 103 cells per well in triplicates in a 96-well plate overnight for attachment, washed, and then fixed for 5 min in 2% formaldehyde and 0.2% glutaraldehyde buffered with PBS. After washing thrice in PBS, the fixed cells were incubated in the staining solution [0.2 mM fluorescein di- β -D-galactopyranoside (FDG, Sigma-Aldrich F2756), 0.2 M citric acid, 0.4 M Na_2_HPO_4_ (citrate–phosphate buffer), 100 mM potassium ferricyanide, 5 M NaCl, 0.2 M MgCl_2_, (all from Sigma Aldrich) in deionized water] in the dark in a humidified incubator at 37 °C for 24 h without CO_2_ supply. Then, an equal volume of the supernatant of each well was transferred to a new 96-well plate for fluorescence measurement. The resultant production of fluorescein was measured using a FluoStar Optima instrument (BMG Labtech) with an excitation of 485 nm and emission at 535 nm. One well containing the reaction mixture without the cells was used as blank for subtracting the background fluorescence.

### Stability assays

Stability in cell culture medium and in mouse and human serum was assayed by adding 625 μL of a 2 mM solution of test compound in PBS (pH 7.4) to 1875 μL of mouse or human serum (Europa Bioproducts, EQSM-0100) pre-warmed at 37 °C or to 1875 μL of cell culture medium. Next, solutions were incubated at 37 °C for 24 h, taking aliquots of 125 μL at different time points (0, 1, 4, 8 and 24 h). Each aliquot was quenched with 500 μL of cold acetonitrile, vortexed, incubated for 10 min in ice and centrifuged at 39000*g* for 10 min. Supernatants were then analyzed by HPLC-MS using SIM mode. Concentrations were quantified by measuring the area under the peak ([M+H]^+^) normalized with an internal standard and converted to the percentage of compound remaining, using the time zero peak area value as 100%.

For measuring the stability in mouse and human liver microsomes, compounds were incubated at 37 °C at a final concentration of 1 or 5 μM in PBS, respectively, together with a solution of nicotinamide adenine dinucleotide phosphate (NADPH) in PBS (final concentration of 2 mM) and a solution of MgCl_2_ in PBS (final concentration of 5 mM). Reactions were initiated by the addition of a suspension of mouse liver microsomes (MLMs) (male CD-1 mice pooled, Sigma-Aldrich) or human liver microsomes (HLMs) (male human pooled, Sigma-Aldrich), respectively, at a final protein concentration of 1 mg/mL. The solutions were vortexed and incubated at 37 °C. Aliquots of 100 μL were quenched at time zero and at seven points ranging to 1 h (MLM) or 2 h (HLM) by pouring into 100 μL of ice-cold acetonitrile. Quenched samples were centrifuged at 10000*g* for 10 min, and the supernatants were filtered through a polytetrafluoroethylene (PTFE) membrane syringe filter (pore size of 0.2 μm, 13 mm of diameter, GE Healthcare Life Sciences). The relative disappearance of the compound under study over the course of the incubation was monitored by HPLC-MS using SIM mode. Concentrations were quantified by measuring the area under the peak ([M+H]^+^) normalized with an internal standard and converted to the percentage of compound remaining, using the time zero peak area value as 100%. The natural logarithm of the remaining percentage versus time data for each compound was fit to a linear regression, and the slope was used to calculate the degradation half-life (t_1/2_). The intrinsic clearance (CL_int_) was calculated by the equation V·ln2/t_1/2_ where V is the incubation volume (in μL) / protein (mg) for microsomal stability.

### Human serum albumin (HSA) binding assay

Determination of the binding of the compound to HSA was performed by incubating a fixed concentration of the compound with different concentrations of immobilized HSA, using the TRANSILXL HSA Binding Kit (TMP-0210-2096, Sovicell). An eightwell unit of the TRANSIL assay plate was used for each compound; six wells contain increasing concentrations of HSA immobilized on silica beads suspended in PBS at pH 7.4, and two wells contain buffer only and serve as references to account for non-specific binding. The TRANSIL assay plate was thawed for 3 h at rt and centrifuged at 750*g* for 5 s. Then, 15 μL of an 80 μM stock solution of the compound in PBS (for a final concentration of 5 μM) was added to each well of the eight-well unit, and the plate was incubated on a plate shaker at 1000 rpm for 12 min at rt. After this time, the plate was centrifuged at 750*g* for 10 min, and 50 μL of the supernatants were transferred for analytical quantification by HPLC-MS using selected ion monitoring (SIM) and quantification was estimated by measuring the area under the peak ([M+H]^+^) normalized with an internal standard. The binding percentage was calculated from the remaining free compound concentration in the supernatant of each well, using the spreadsheet and algorithms supplied with the kit.

### Animal experiments

All scientific procedures with animals were conformed to EU Directive 2010/63EU and Recommendation 2007/526/EC, enforced in Spanish law under Real Decreto 53/2013. Animal protocols were approved by the Committee of Animal Experimentation of Universidad Complutense de Madrid and the Animal Protection Area of the Comunidad Autónoma de Madrid (PROEX 159/18). Animal studies were carried out in *Lmna*^*G609G/G609G*^ knock-in mice ubiquitously expressing progerin^20^ and control *Lmna*^+/+^ littermates. Mice were maintained in the animal facility of the Universidad Complutense de Madrid under specific pathogen free conditions. For pharmacokinetic studies, compound **21** was administered intraperitoneally and blood was collected at the selected time points post-dose (n=3 per time point) by cardiac puncture. Blood was allowed to clot at room temperature for 30 min and centrifuged at 4 °C for 10 min at 16000*g*. The supernatant was transferred to a clean polypropylene tube and stored at −80 °C until analysis. For analysis, a volume of cold acetonitrile was added to the serum. The sample was incubated in an ice bath for 10 min and centrifuged at 4 °C for 10 min at 16000*g*. The resulting organic layer was filtered through a polytetrafluoroethylene filter (0.2 μm, 13 mm diameter, Fisher Scientific) and 20 μL of the sample analysed by LC-MS/MS at the UCM’s Mass Spectrometry CAI. Separation was performed using a Phenomenex Gemini 5 μm C18 110A 150×2 mm column (run time 10 min; flow 0.5 mL/min; gradient: 5 min 5% B – 3 min 100% B – 2 min 5% B; Phase A: water with formic acid 0.1%; Phase B: acetonitrile). The entire LC eluent was directly introduced to an electrospray ionization (ESI) source operating in the positive ion mode for LC MS/MS analysis on a Shimadzu LCMS8030 triple quadrupole mass spectrometer coupled to UHPLC with an oven temperature of 31.5 °C. The mass spectrometer ion optics were set in the multiple reaction monitoring mode and the transition selected for quantification was 382.10 > 217.15 (CE: −19 V).

For in vivo treatment, compound **21** was administrated at a concentration of 40 mg/kg diluted in sterile corn oil (Sigma-Aldrich) intraperitoneally five times per week starting at the age of 6 weeks. The same quantity of corn oil was injected to control *Lmna*^*G609G/G609G*^ and *Lmna*^*+/+*^ mice. Treatments did not produce any apparent damage or stress in mice. The weight of each mouse was registered once a week during the duration of the treatment.

To analyze blood glucose, animals were starved for 7 h and glucose levels were measured with a FreeStyle Optium Neo glucometer (Abbott) using blood from the tail vein.

Forepaw strength of in the three groups of mice was measured as previously described.^33^ Briefly, seven weights of 20, 33, 46, 59, 72, 85 and 98 grams were used. The ability of each mouse to hold each weight from lighter to heavier was determined. A final total score is calculated as the product of the number of links in the heaviest chain held for the full 3 seconds, multiplied by the seconds it is held. If the heaviest weight is dropped before 3 seconds, an intermediate value is calculated.

In vivo magnetic resonance imaging (MRI) was performed at the Preclinical MRI Unit of the Complutense University using a 1-Tesla benchtop MRI scanner (Icon 1T-MRI; Bruker BioSpin GmbH, Ettlingen, Germany). MRI system consists of a 1 T permanent magnet (without extra cooling required for the magnet) with a gradient coil that provides a gradient strength of 450 mT/m. Animal monitoring systems and solenoid radiofrecuency coil (oval shape, inner diameter 59 × 50 mm^2^ and length 90 mm) are integrated into the magnet bed. They allowed both the animals accurate positioning and the correct monitoring of vital signs during acquisition. Animals were anesthetized using 2% isofluorane (IsoFlo, Zoetis, NJ, USA) and they were positioned unrestrained in order to visualize the natural curvature of their spinal corn. The main MRI experiment consisted of three dimensional images (0.3 × 0.3 × 1.0 mm^3^). It was acquired using a gradient echo sequence with a repetition time of 160 ms, an echo time of 3.5 ms, and a flip angle of 45°. The total acquisition time was less than 6 min. All MRI images were analysed using ImageJ software.

### Pathological analysis and immunofluorescence

After 6 weeks of treatment, 12-week-old mice were euthanized by CO_2_ inhalation. Immediately after sacrifice, the thoracic and abdominal cavities were opened. Mice were perfused with cold PBS and spleen, thymus, heart, and aorta were excised and used for different protocols. For immunohistofluorescence, heart and aorta (aortic arch and thoracic aorta) segments were fixed overnight in 4% paraformaldehyde and included in paraffin to prepare 4‐μM sections using a microtome. Antigen retrieval was performed with 10 mM sodium citrate buffer (pH 6). Samples were blocked and permeabilized for 1 h at rt in PBS containing 0.3% TritonX‐100 (9002‐93‐1, Sigma), 5% normal goat serum (005‐000‐001, JacksonImmunoResearch), and 5% bovine serum albumin (BSA, A7906, Sigma). Sections were incubated overnight at 4°C with rabbit anti‐progerin polyclonal antibody (1/300) generated using peptide immunogens and standard immunization procedures (S.Nourshargh et al., manuscript in preparation) diluted in PBS containing 2.5% normal goat serum. Secondary antibody (A‐21245, goat anti‐rabbit Alexa Fluor 647), and Hoechst 33342 stain (B2261, Sigma) were diluted in 2.5% normal goat serum in PBS and mounted using Fluoromount G imaging medium (00‐4958‐02, Affymetrix eBioscience). Fluorescence images were acquired with a Zeiss LSM 700 confocal microscope. Image analysis was performed using ImageJ Fiji software by a researcher blinded to genotype.

### Statistics

All data were analyzed using GraphPad Prism 6 software. Data were presented as mean±standard error of the media (SEM) with at least three biologically independent experiments. Representative morphological images were taken from at least three biologically independent experiments with similar results. The Student t test or one-way ANOVA were used for comparison between groups. Survival analysis was performed using the Kaplan−Meier method. Differences between survival distributions were analyzed using the log rank test. Hazard ratio and confidence interval were obtained by Mantel−Haenszel analysis. Differences with P < 0.05 were considered significant.

## Data availability

The data that support the findings of this study are available from the corresponding author upon reasonable request.

---

**Extended data Fig. 1.**
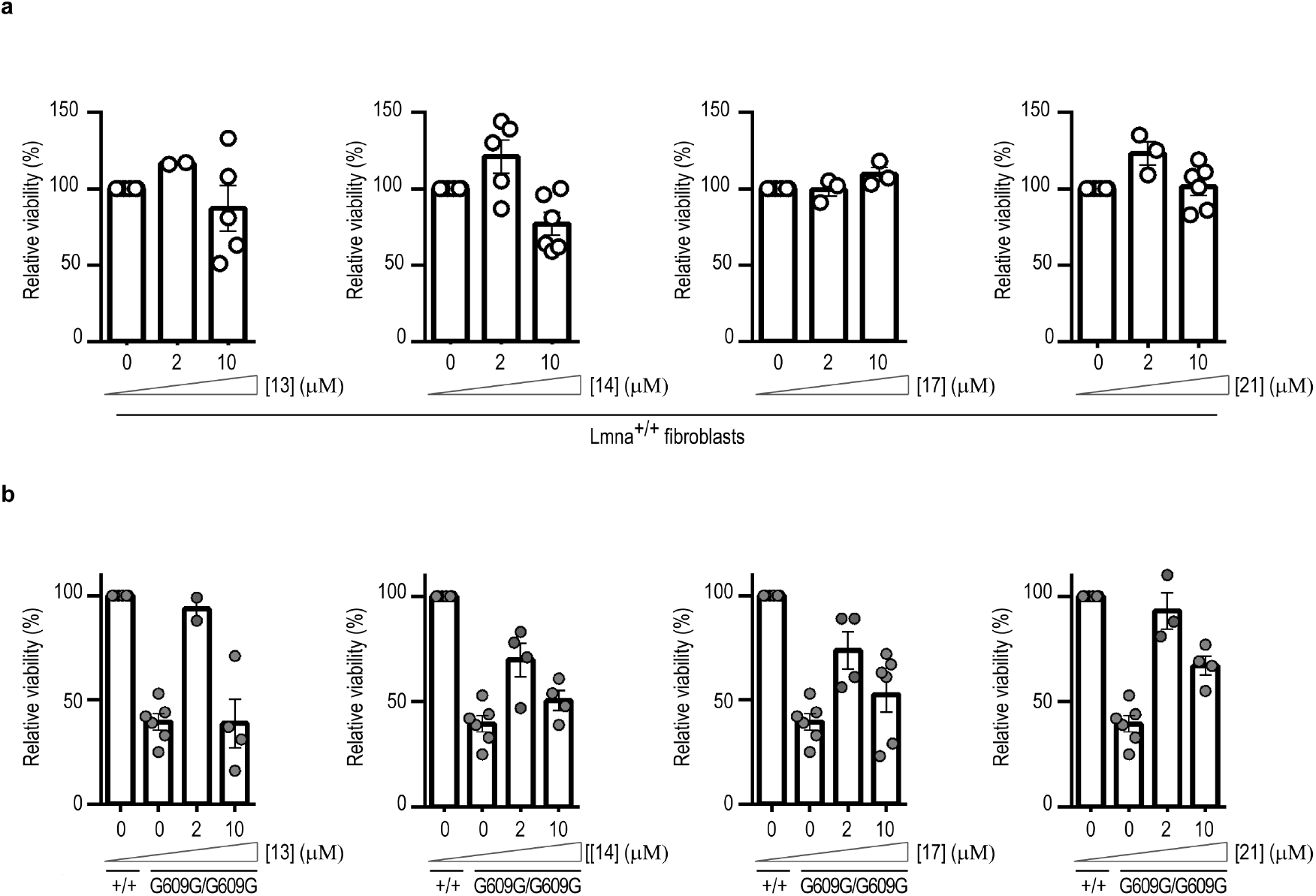
Effect of compounds in the viability of *Lmna*^*+/+*^ or *Lmna*^*G609G/G609G*^ mouse fibroblasts. **a**, *Lmna*^*+/+*^ or **b**, *Lmna*^*G609G/G609G*^ cells were incubated with 0.1% DMSO (vehicle, veh) or different compounds at 2 μM or 10 μM during 3 days and viability was determined in MTT assay. Plots show percentage of cell viability (n≥3 independent experiments).

**Extended data Fig. 2.**
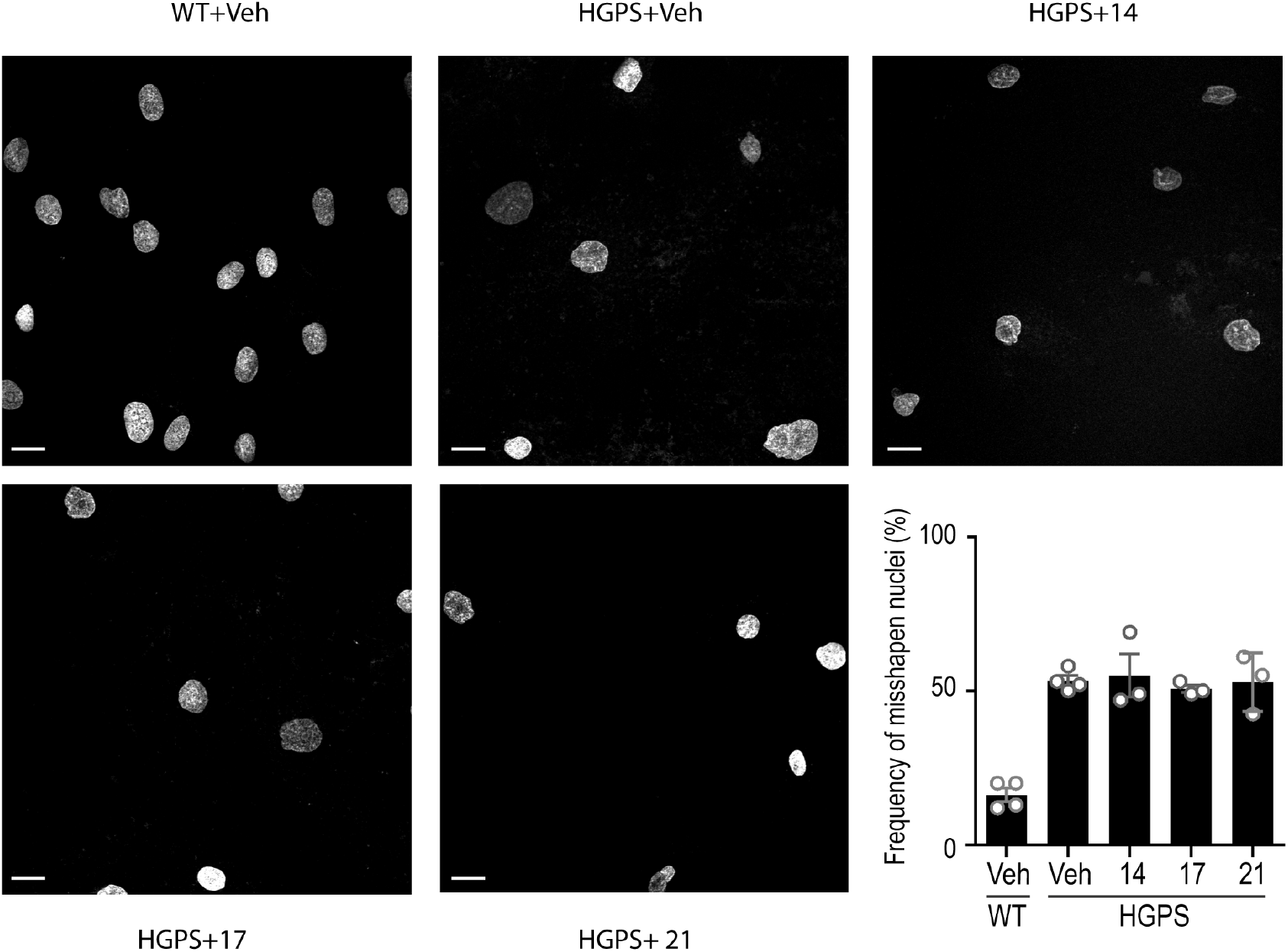
ICMT inhibitors do not affect misshapen nuclei HGPS human cells. Immunofluorescence images of wild type (WT) or HPGS human fibroblasts treated with 0.1% DMSO (vehicle, veh) or different ICMT inhibitors at 2 μM during 17 days and stained with Hoechst. Plot shows percentage average of misshapen nuclei in at least five images per condition from ≥3 experiments. No significant differences (one way ANOVA) were found between treated and non-treated HGPS cells. Scale bar: 20 μm.

**Extended data Fig. 3.**
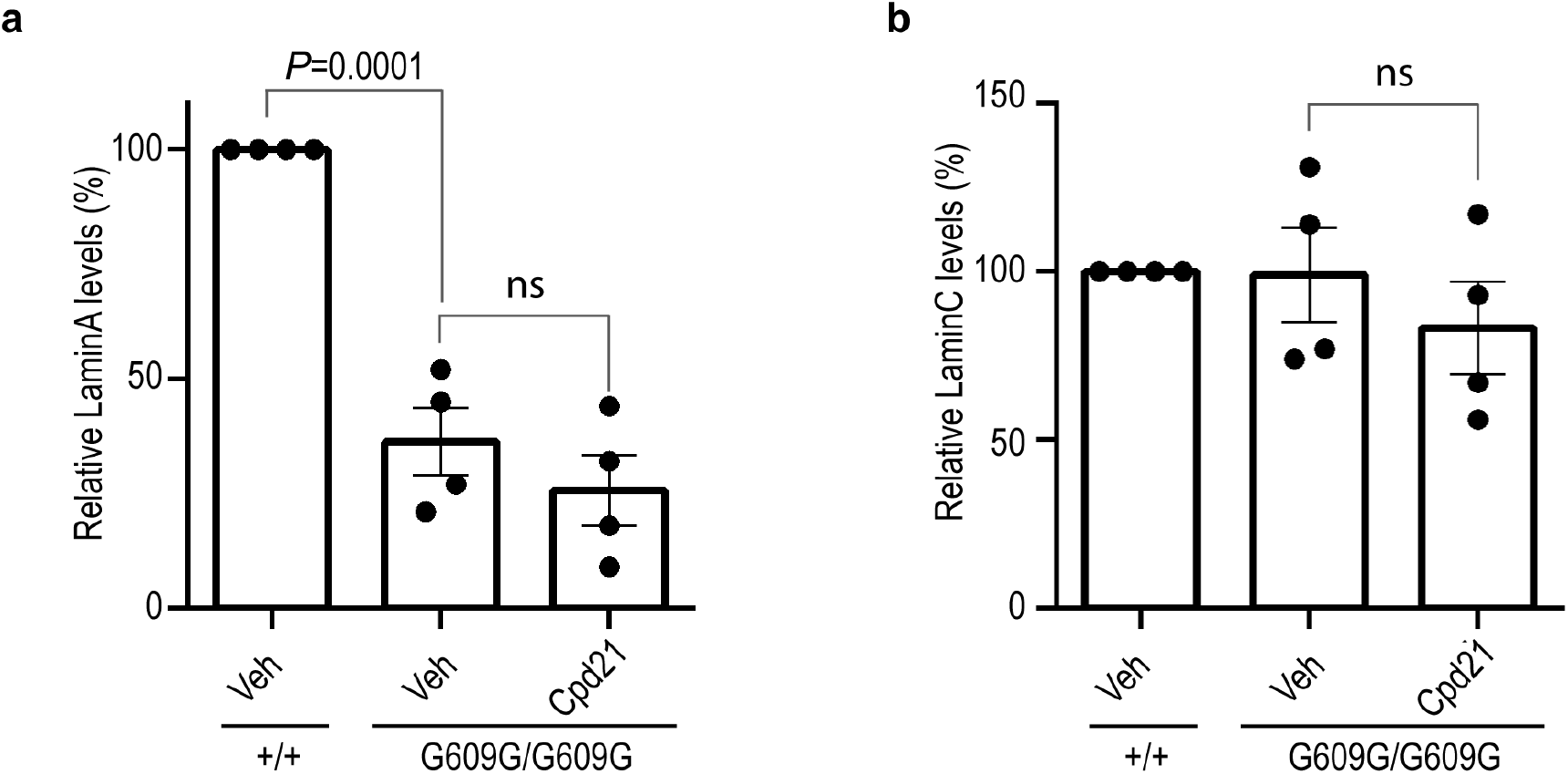
ICMT inhibitors do not affect the levels or lamin A or lamin C. *Lmna*^*+/+*^ or *Lmna*^*G609G/G609G*^ fibroblasts were incubated with 0.1% DMSO (vehicle, veh) or compound **21** at 2 μM during 14 days and lamin A (a) or lamin C (b) levels were analysed by western blot (n=4 independent experiments; plots show protein level±sem normalized against GAPDH levels, two-tailed Student’s t-test).

**Extended data Fig. 4.**
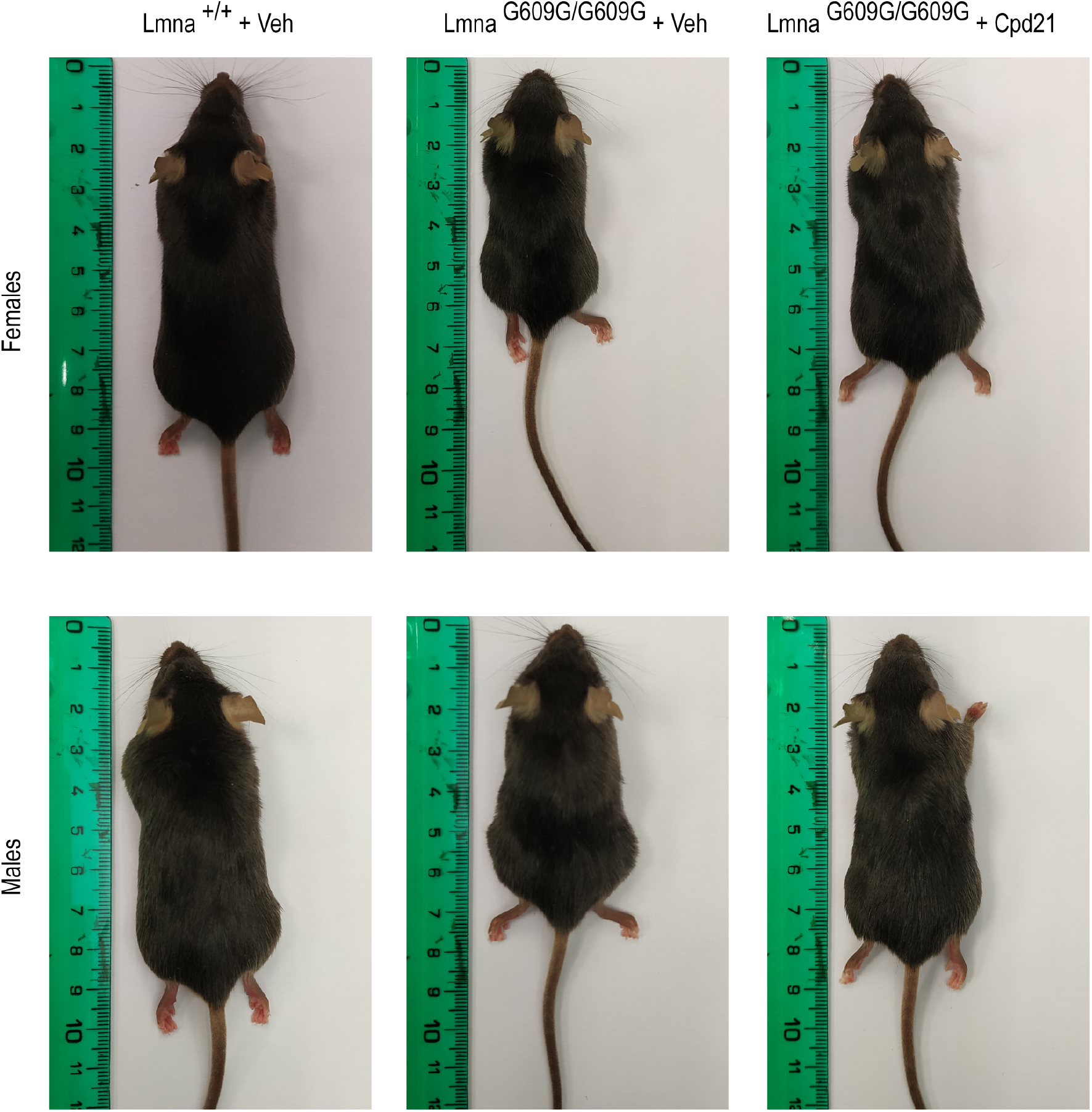
Compound 21 improved size in males and females. Photographs of three months old sex-matched mice *Lmna*^*+/+*^ or *Lmna*^*G609G/G609G*^ treated with vehicle compared to *Lmna*^*G609G/G609G*^ mice treated with compound **21**.

**Extended data Fig. 5.**
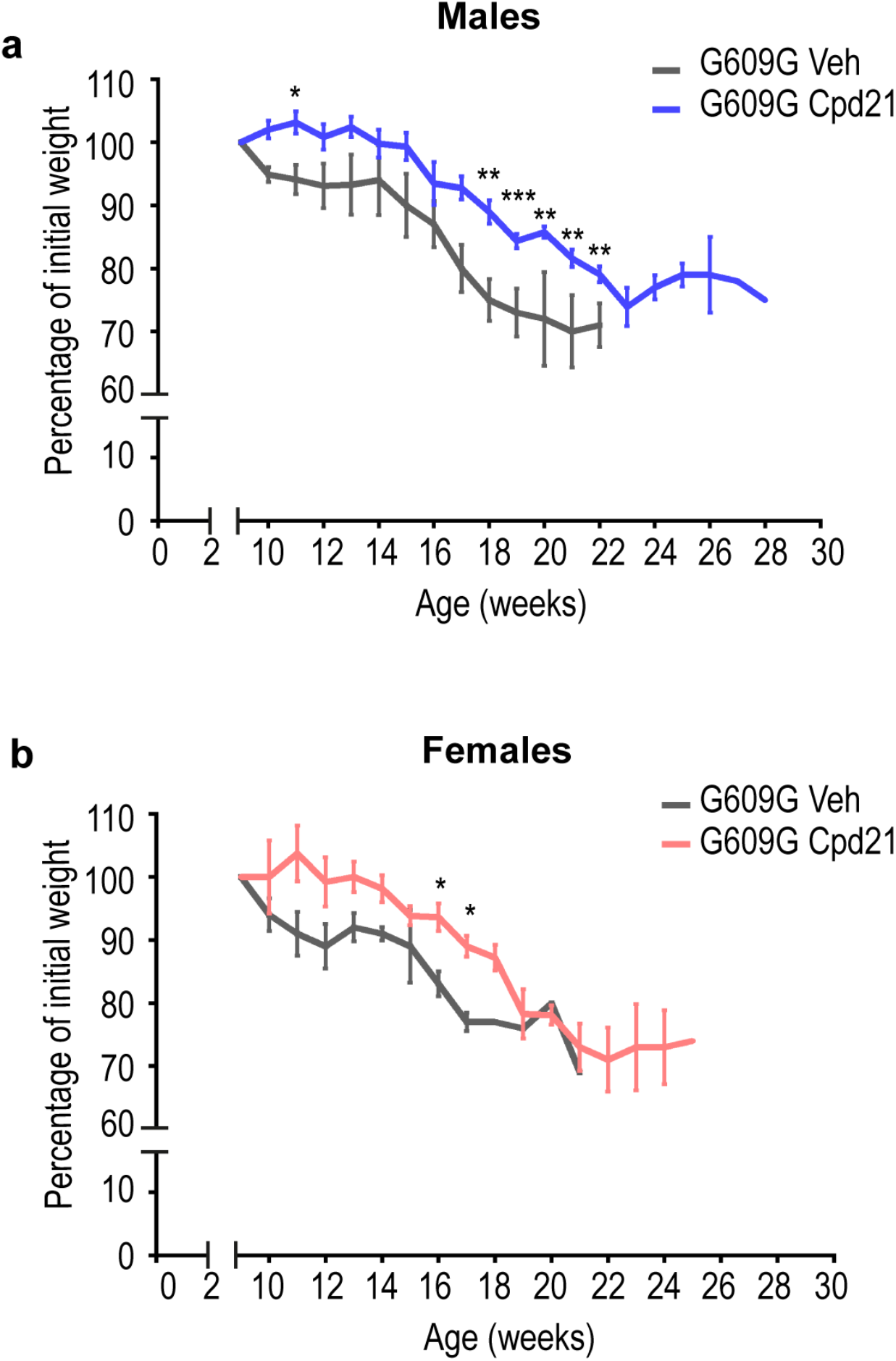
Progression of body weight of male or female *Lmna*^*G609G/G609G*^ mice treated with compound 21 or with vehicle. **a**, Body weight versus age plot of untreated (n=4) and treated (n=6) male *Lmna*^*G609G/G609G*^ mice (plot shows mean±sem, two-tailed Student’s t-test, p-value *<0.05, **<0.005, ***<0.0005). **b**, Body weight versus age plot of untreated (n=2) and treated (n=4) female *Lmna*^*G609G/G609G*^ mice (plot shows mean±sem, two-tailed Student’s t-test, p-value *<0.05, **<0.005, ***<0.0005).

**Extended data Fig. 6.**
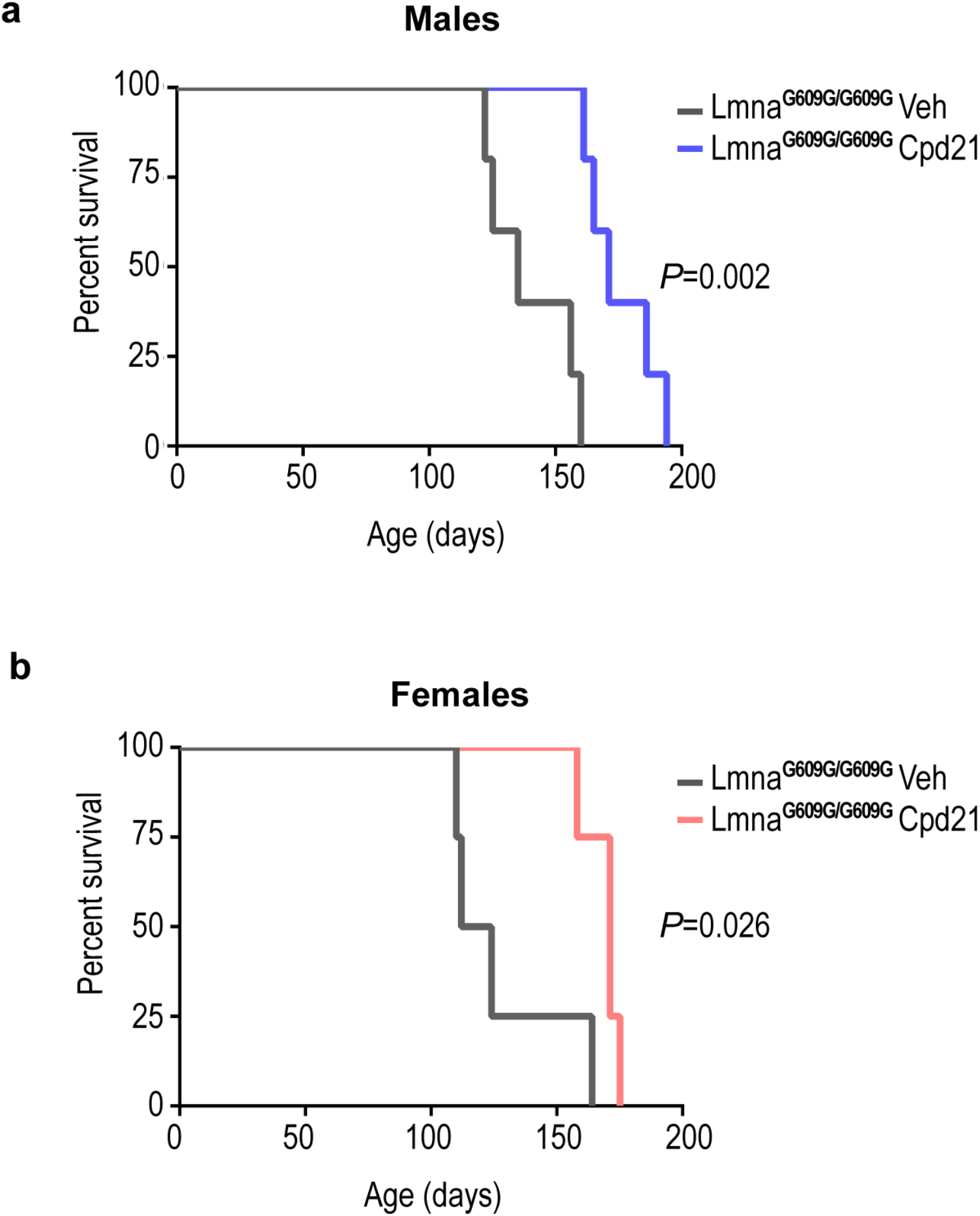
Survival of male or female *Lmna*^*G609G/G609G*^ mice treated with compound 21 or with vehicle. **a**, Kaplan-Meier survival plot of *Lmna*^*G609G/G609G*^ male mice treated with compound **21** (n=6) or vehicle (n=5), (P=0.002, log-rank/Mantel-Cox test). **b**, Kaplan-Meier survival plot of *Lmna*^*G609G/G609G*^ female mice treated with compound **21** (n=4) or vehicle (n=4), (P=0.026, log-rank/Mantel-Cox test).

**Extended data Fig. 7.**
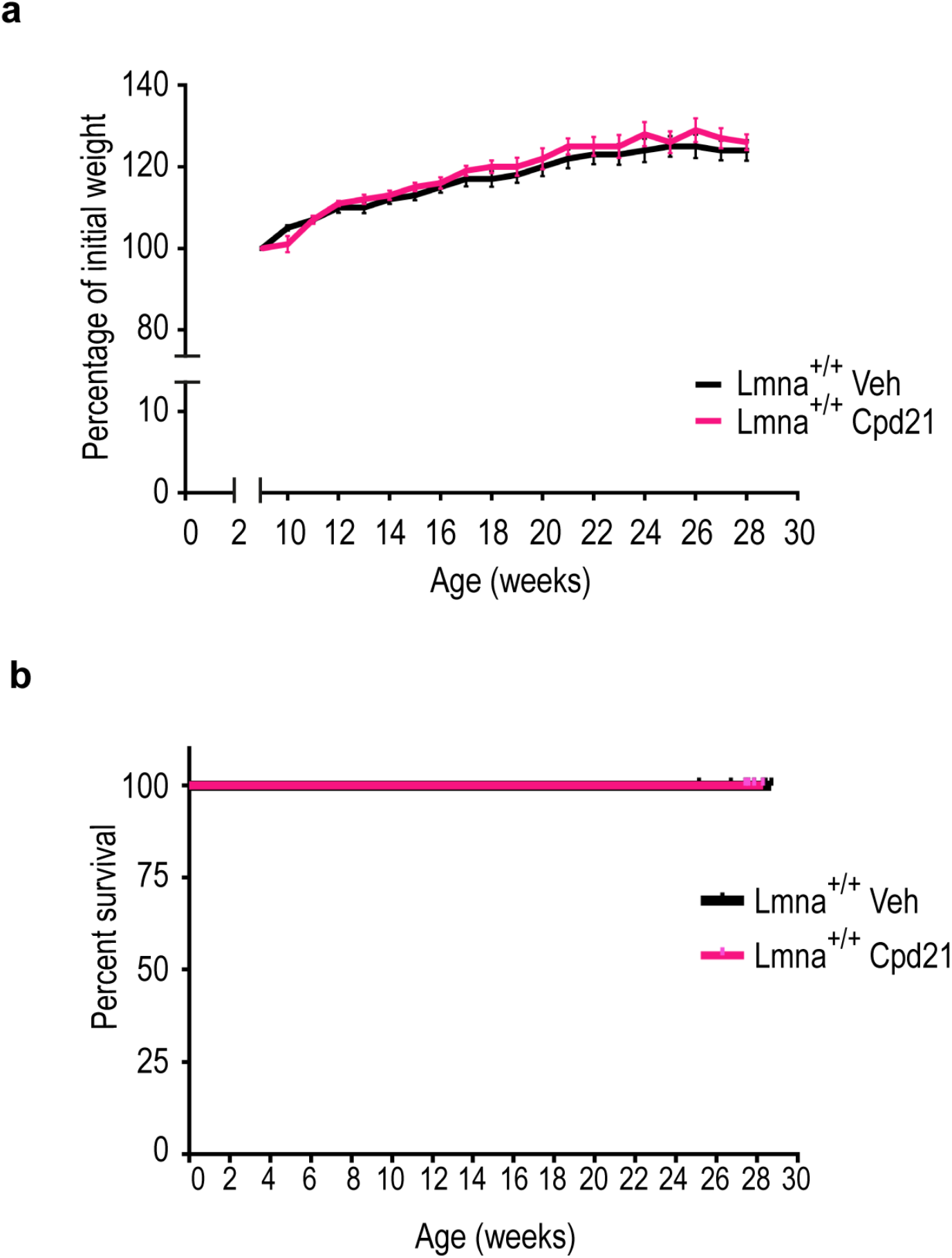
Progression of body weight and survival of wild type (WT) mice treated with compound 21 or with vehicle. **a**, Plot of body weight versus age of *Lmna*^*+/+*^ mice treated with compound **21** (n=7) or with vehicle (n=10) (plot shows mean±sem). **b**, Kaplan-Meier survival plot of *Lmna*^*+/+*^ mice treated with compound **21** (n=7) or vehicle (n=10).

**Extended data Fig. 8.**
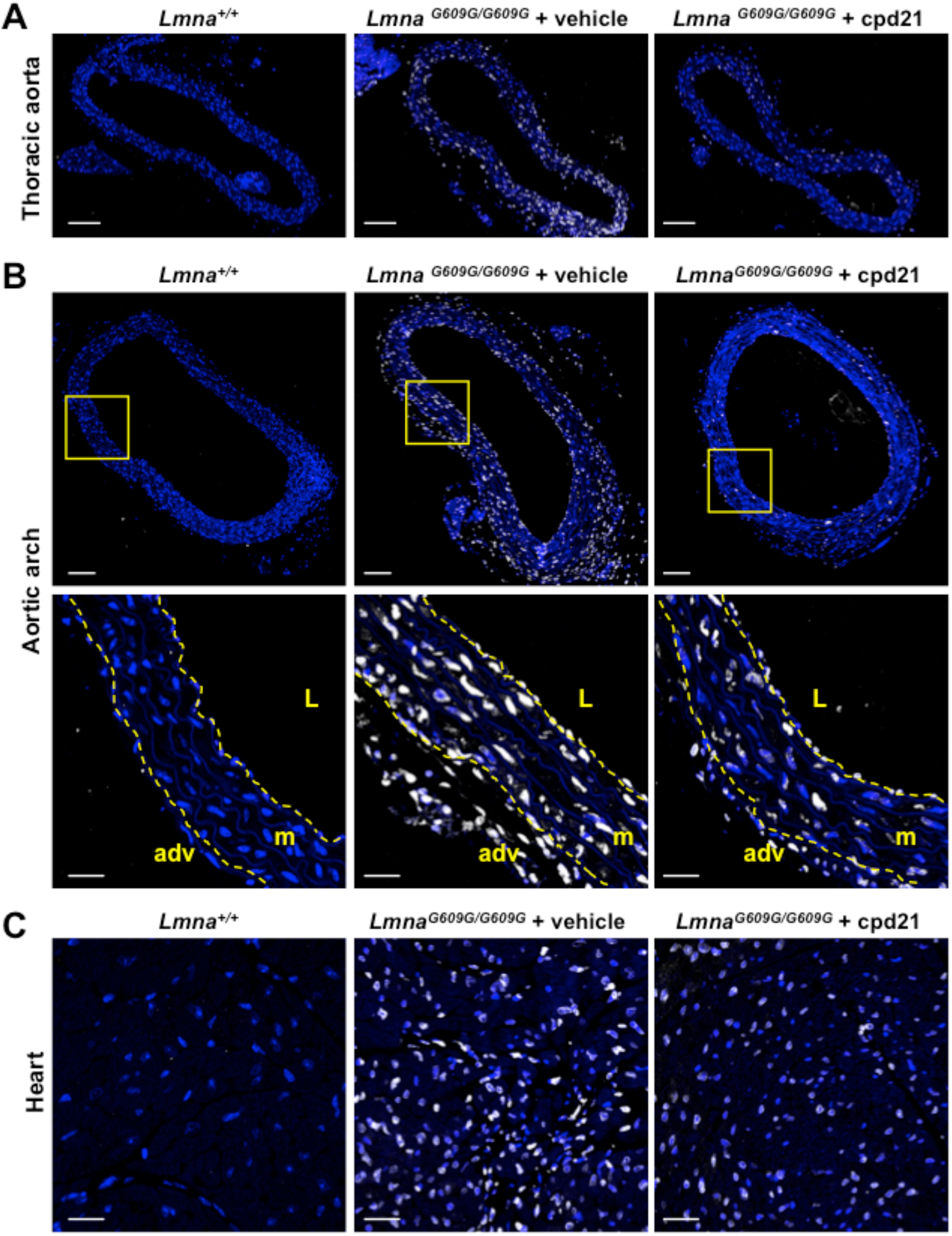
Reduced progerin expression in progeric *Lmna* ^*G609G/G609G*^ mice treated with compound 21. Mice of the indicated genotypes were treated with compound **21** or vehicle, starting at 6 weeks of age, and were sacrificed at 12 weeks of age. Cross-sections of thoracic aorta, aortic arch and heart were stained with anti‐progerin antibody (white) and Hoechst 33342 to visualize nuclei (blue). Representative immunofluorescence images are shown. **A**. Thoracic aorta. Scale bar: 100 μm. **B**. Aortic arch. Scale bar: 100 μm (entire image) and 20 μm (magnified view). L, lumen; m, media; adv, adventitia. **C**. Myocardial tissue. Scale bar: 20 μm.

