## Supplementary Information for "Isoprenylcysteine carboxylmethyltransferase-based therapy for Hutchinson–Gilford progeria syndrome"

Supporting Table 1

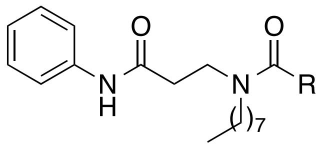


**2-25**

| **Cpd** | **R** | **ICMT inhibition (%)**[a] | **Viability (%)**[b] | |
|---|---|---|---|---|
| | | | Wild type mouse fibroblasts | Progeroid mouse fibroblasts |
| **Vehicle** | | 0 | 100 | 41±3 |
| **2** | 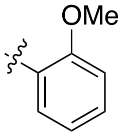 | 32 | - | - |
| **3** | 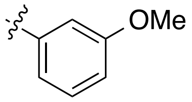 | 26 | - | - |
| **4** | 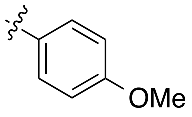 | 52 | - | - |
| **5** | 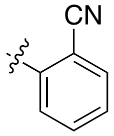 | 47 | | |
| **6** | 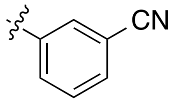 | 53 | | |
| **7** | 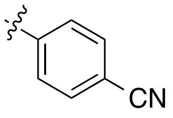 | 66 | 86±2 | 56±4 |
| **8** | 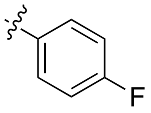 | 42 | - | - |
| **9** | 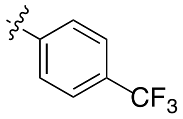 | 18 | - | - |

| | | | | |
|---|---|---|---|---|
| **10** | 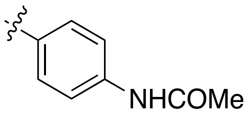 | 10 | - | - |
| **11** | 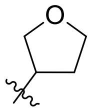 | 41 | - | - |
| **12** | 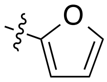 | 49 | - | - |
| **13** | 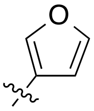 | 81 | 116±2 | 77±5 |
| **14** | 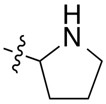 | 62 | 121±11 | 70±8 |
| **(*R*)-14** | 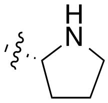 | 61 | 114±6 | 80±7 |
| **(*S*)-14** | 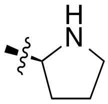 | 64 | 114±10 | 70±5 |
| **15** | 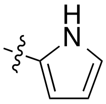 | 75 | 107±5 | 45±6 |
| **16** | 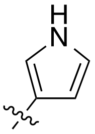 | 81 | 109±4 | 53±13 |
| **17** | 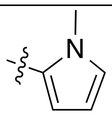 | 75 | 99±5 | 74±9 |
| **18** | 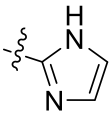 | 34 | - | - |
| **19** | 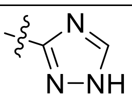 | 0 | - | - |
| **20** | 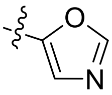 | 56 | - | - |
| **21** | 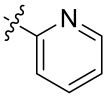 | 73 | 113±9 | 93±9 |

| | | | | |
|---|---|---|---|---|
| **22** | 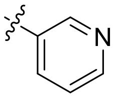 | 38 | - | - |
| **23** | 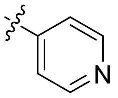 | 30 | - | - |
| **24** | 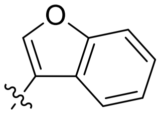 | 47 | - | - |
| **25** | 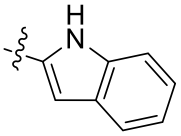 | 15 | - | - |

[a]ICMT inhibition values were determined at a concentration of 50 μM. Results are expressed as the mean from at least two independent experiments performed in triplicate. The sem is within a 10% of the mean value. [b]Cell viability was determined by the MTT assay. Data are expressed as the mean±sem from three independent experiments performed in triplicate at a compound concentration of 2 μM. The percentage of viability is expressed considering 100% viability in vehicle-treated wild type fibroblasts.

### 3. Synthesis of compounds 1-25

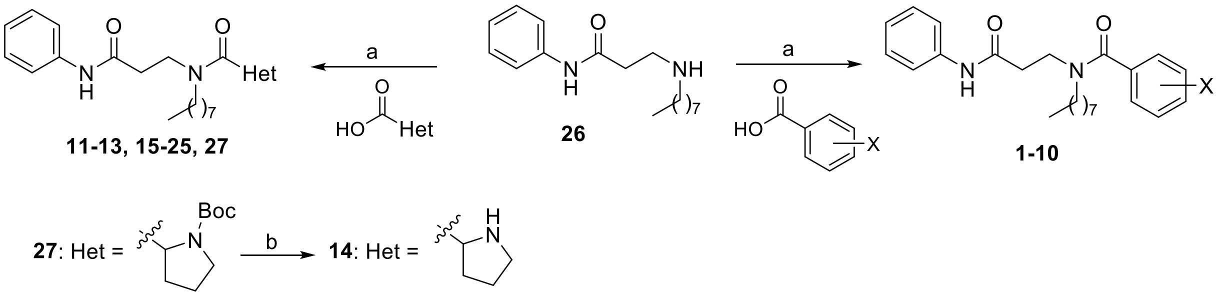


**Scheme S1.** Reagents and conditions: a) EDC, HOBT, DCM, rt, 16 h, 36-98%; b) TFA, DCM, rt, 70-90%.

$N^3$-Octyl-$N^1$-phenyl-β-alaninamide (**26**) was synthesized as previously described and its spectroscopic data are in agreement with those reported.[1]

---

[1] Marín-Ramos NI, Balabasquer M, Ortega-Nogales FJ, Torrecillas IR, Gil-Ordóñez A, Marcos-Ramiro B, Aguilar-Garrido P, Cushman I, Romero A, Medrano FJ, Gajate C, Mollinedo F, Philips MR, Campillo M, Gallardo M, Martín-Fontecha M, López-Rodríguez ML, Ortega-Gutiérrez S. *J. Med. Chem.* **2019**, *11*. 6035-6046. doi: 10.1021/acs.jmedchem.9b00145

·**General procedure for the synthesis of compounds 1-13, 15-25 and 27.** To a solution of the corresponding carboxylic acid (1-2 equiv) in anhydrous DCM (4 mL/mmol), EDC (1-2 equiv) and HOBt (1-2 equiv) were added. The reaction mixture was stirred at rt for 1 h. Then, a solution of the secondary amine **26** (1 equiv) in anhydrous DCM (2 mL/mmol) was added and the reaction mixture was stirred at rt for 16 h. The reaction crude was washed with saturated aqueous solutions of $NaHCO_3$ and NaCl, consecutively. The organic extracts were dried over $Na_2SO_4$, filtered, and the solvent removed under reduced pressure. The residue was purified by column chromatography obtaining the desired amides.

***N*$^2$-Benzoyl-*N*$^2$-octyl-*N*$^1$-phenyl-β-alaninamide (1)**. Obtained from amine **26** (150 mg, 0.54 mmol), benzoic acid (132 mg, 1.1 mmol), EDC (207 mg, 1.1 mmol) and HOBt (146 mg, 1.1 mmol) in 91% yield (164 mg). Chromatography: hexane/EtOAc, 9:1.

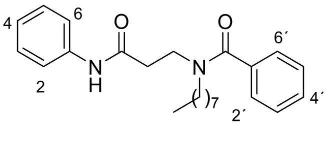


$R_f$ (hexane/EtOAc, 7:3): 0.28. Mp: 65-67 °C. IR (ATR, ν): 3307 (NH), 1686 (CON), 1607, 1546, 1498, 1439 (Ar). $^1$H-NMR ($CDCl_3$, δ): 0.85 (t, *J* = 6.9 Hz, 3H, $CH_3$), 1.09-1.15 (m, 10H, $(CH_2)_5CH_3$), 1.50 (m, 2H, $CH_2(CH_2)_5CH_3$), 2.75 (m, 2H, $CH_2CO$), 3.25 (m, 2H, $(CH_2)_6CH_2N$), 3.81 (m, 2H, $COCH_2CH_2N$), 7.04 (t, *J* = 7.4 Hz, 1H, $H_4$), 7.21-7.27 (m, 2H, $H_{3'}$, $H_{5'}$), 7.32-7.40 (m, 5H, $H_3$, $H_5$, $H_{2'}$, $H_{4'}$, $H_{6'}$), 7.51 (d, *J* = 7.7 Hz, 2H, $H_2$, $H_6$), 9.35 (br s, 1H, NH). $^{13}$C-NMR ($CDCl_3$, δ): 14.0 ($CH_3$), 22.6, 26.4, 28.8, 29.0 (2C), 31.7 ($(CH_2)_6CH_3$), 36.2 ($CH_2CO$), 42.6 ($CH_2N$), 50.6 ($(CH_2)_6CH_2N$), 119.9 ($C_2$, $C_6$), 123.9 ($C_4$), 126.4 ($C_{2'}$, $C_{6'}$), 128.5 ($C_{3'}$, $C_{5'}$), 128.8 ($C_3$, $C_5$), 129.5 ($C_{4'}$), 136.4 ($C_{1'}$), 138.6 ($C_1$), 169.8, 172.7 (2CO). MS (ESI, m/z): 381.2 $[M+H]^+$. Elemental analysis calculated for $C_{24}H_{32}N_2O_2$: C, 75.75; H, 8.48; N, 7.36; found: C, 75.45; H, 8.21; N, 7.41.

***N*$^2$-(2-Methoxyphenylcarbonyl)-*N*$^2$-octyl-*N*$^1$-phenyl-β-alaninamide (2).** Obtained from amine **26** (100 mg, 0.36 mmol), 2-methoxybenzoic acid (66 mg, 0.43 mmol), EDC (67 mg, 0.43 mmol) and HOBt (58 mg, 0.43 mmol) in 74% yield (110 mg). Chromatography: hexane/EtOAc, 1:1.

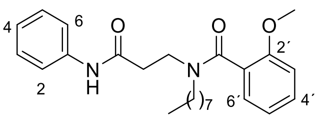


$R_f$ (hexane/EtOAc, 1:1): 0.42. IR (ATR, ν): 3270 (NH), 1686, 1547 (CON), 1599, 1498, 1441 (Ar). $^1$H-NMR ($CDCl_3$, δ): amide rotamers A:B, 9:1; 0.86 (t, *J* = 7.0 Hz, 3H, $CH_3CH_2$), 1.08-1.31 (m, 10H, $(CH_2)_5CH_3$), 1.44-1.51 (m, 2H, $CH_2(CH_2)_5CH_3$ rotamer A), 1.52-1.73 (m, 2H, $CH_2(CH_2)_5CH_3$ rotamer B), 2.17 (m, 2H, $CH_2CO$ rotamer B), 2.39-2.48 (m, 2H, $(CH_2)_6CH_2N$ rotamer B), 2.80-2.85 (m, 2H, $CH_2CO$ rotamer A), 3.13-3.20 (m, 2H, $(CH_2)_6CH_2N$ rotamer A), 3.42-3.57 (m, 2H, $COCH_2CH_2N$ rotamer B), 3.67 (s, 3H, $CH_3O$ rotamer A), 3.72-4.06 (m, 5H, $COCH_2CH_2N$ rotamer A, $CH_3O$ rotamer B), 6.86 (d, *J* = 8.3 Hz, 1H, $H_{3'}$), 6.95 (t, *J* = 7.4 Hz, 1H, $H_{5'}$), 7.04 (t, *J* = 7.4 Hz, 1H, $H_4$), 7.19 (dd, *J* = 7.4, 1.7 Hz, 1H, $H_{6'}$), 7.24 (t, *J* = 8.1 Hz, 2H, $H_3$, $H_5$), 7.32 (td, *J* = 7.1, 1.7 Hz, 1H, $H_{4'}$), 7.53 (d, *J* = 7.7 Hz, 2H, $H_2$, $H_6$), 9.47 (br s, 1H, NH). $^{13}$C-NMR ($CDCl_3$, δ): 14.2 ($CH_3CH_2$), 22.7, 26.5, 28.6, 29.1 (2C), 31.8 ($(CH_2)_6CH_3$), 36.3 ($CH_2CO$), 42.0 ($CH_2N$), 49.9 ($(CH_2)_6CH_2N$), 55.4 ($CH_3O$), 110.9 ($C_{3'}$), 120.0 ($C_2$, $C_6$), 120.8 ($C_{5'}$), 123.7 ($C_4$), 125.9 ($C_{1'}$), 127.6 ($C_{6'}$), 128.7 ($C_3$, $C_5$), 130.5 ($C_{4'}$), 138.8 ($C_1$), 155.2 ($C_{2'}$), 170.0, 170.5 (2CO). HRMS (ESI, m/z): Calculated for $C_{25}H_{34}N_2O_3Na$ $[M+Na]^+$: 433.2467; found: 433.2466.

***N*$^2$-(3-Methoxyphenylcarbonyl)-*N*$^2$-octyl-*N*$^1$-phenyl-β-alaninamide (3).** Obtained from amine **26** (100 mg, 0.36 mmol), 3-methoxybenzoic acid (66 mg, 0.43 mmol), EDC (67 mg, 0.43 mmol) and HOBt (58 mg, 0.43 mmol) in 78% yield (115 mg). Chromatography: hexane/EtOAc, 1:1.

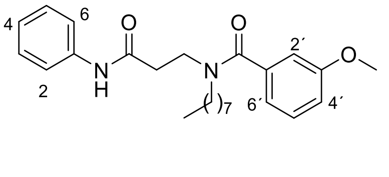


$R_f$ (hexane/EtOAc, 1:1): 0.48. IR (ATR, ν): 3273 (NH), 1687, 1547 (CON), 1601, 1498, 1442 (Ar). $^1$H-NMR ($CDCl_3$, δ): amide rotamers A:B, 9:1; 0.86 (t, *J* = 6.9 Hz, 3H, C*H*$_3$CH$_2$), 1.12-1.27 (m, 10H, (C*H*$_2$)$_5$CH$_3$), 1.52 (m, 2H, C*H*$_2$(CH$_2$)$_5$CH$_3$), 2.48 (m, 2H, CH$_2$CO rotamer B), 2.77 (t, *J* = 6.0 Hz, 2H, CH$_2$CO rotamer A), 3.27 (t, *J* = 7.5 Hz, 2H, (CH$_2$)$_6$C*H*$_2$N rotamer A), 3.45 (m, 2H, (CH$_2$)$_6$C*H*$_2$N rotamer B), 3.69 (s, 3H, CH$_3$O), 3.81-3.92 (m, 2H, COCH$_2$C*H*$_2$N), 6.83 (s, 1H, $H_{2'}$), 6.90 (t, *J* = 7.4 Hz, 1H, $H_{5'}$), 6.91 (d, *J* = 8.6 Hz, 1H, $H_{4'}$), 7.06 (t, *J* = 7.4 Hz, 1H, $H_4$), 7.23-7.29 (m, 3H, $H_3$, $H_5$, $H_{6'}$), 7.54 (d, *J* = 7.9 Hz, 2H, $H_2$, $H_6$), 9.39 (br s, 1H, NH). $^{13}$C-NMR (CDCl$_3$, δ): 14.2 (*C*H$_3$CH$_2$), 22.7, 26.5, 28.9, 29.1, 29.2, 31.8 ((*C*H$_2$)$_6$CH$_3$), 36.2 (*C*H$_2$CO), 42.7 (CH$_2$N), 50.7 ((CH$_2$)$_6$*C*H$_2$N), 55.3 (CH$_3$O), 111.7 ($C_{2'}$), 115.6 ($C_{6'}$), 118.6 ($C_{5'}$), 120.0 ($C_2$, $C_6$), 124.0 ($C_4$), 128.9 ($C_3$, $C_5$), 129.8 ($C_{4'}$), 137.6 ($C_{1'}$), 138.6 ($C_1$), 159.7 ($C_{3'}$), 169.9, 172.6 (2CO). HRMS (ESI, m/z): Calculated for $C_{25}H_{33}N_2O_3$ [M-H]$^-$: 409.2496; found: 409.2491.

***N*$^2$-(4-Methoxyphenylcarbonyl)-*N*$^2$-octyl-*N*$^1$-phenyl-β-alaninamide (4).** Obtained from amine **26** (50 mg, 0.18 mmol), 4-methoxybenzoic acid (33 mg, 0.21 mmol), EDC (34 mg, 0.21 mmol) and HOBt (29 mg, 0.21 mmol) in 73% yield (52 mg). Chromatography: hexane/EtOAc, 7:3 to 3:7.

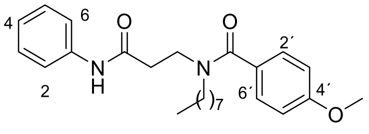


*R*$_f$ (hexane/EtOAc, 7:3): 0.31. IR (ATR, ν): 3309 (NH), 1688, 1547 (CON), 1607, 1504, 1441 (Ar). $^1$H-NMR (CDCl$_3$, δ): 0.86 (t, *J* = 6.9 Hz, 3H, C*H*$_3$CH$_2$), 1.14-1.26 (m, 10H, (C*H*$_2$)$_5$CH$_3$), 1.53 (m, 2H, C*H*$_2$(CH$_2$)$_5$CH$_3$), 2.77 (m, 2H, CH$_2$CO), 3.32 (m, 2H, (CH$_2$)$_6$C*H*$_2$N), 3.81-3.86 (m, 5H, COCH$_2$C*H*$_2$N, CH$_3$O), 6.86 (d, *J* = 8.8 Hz, 2H, $H_{3'}$, $H_{5'}$), 7.07 (t, *J* = 7.4 Hz, 1H, $H_4$), 7.29 (t, *J* = 9.2 Hz, 2H, $H_3$, $H_5$), 7.30 (m, 2H, $H_{2'}$, $H_{6'}$), 7.56 (d, *J* = 7.7 Hz, 2H, $H_2$, $H_6$), 9.31 (br s, 1H, NH). $^{13}$C-NMR (CDCl$_3$, δ): 14.2 (*C*H$_3$CH$_2$), 22.7, 26.6, 28.9, 29.2 (2C), 31.8 ((*C*H$_2$)$_6$CH$_3$), 36.4 (*C*H$_2$CO), 42.8 (CH$_2$N), 50.9 ((CH$_2$)$_6$*C*H$_2$N), 55.4 (CH$_3$O), 113.8 ($C_{3'}$, $C_{5'}$), 120.0 ($C_2$, $C_6$), 124.0 ($C_4$), 128.5 ($C_3$, $C_5$), 128.9 ($C_{2'}$, $C_{6'}$), 138.6 ($C_1$), 143.0 ($C_{1'}$), 160.7 ($C_{4'}$), 169.9, 172.9 (2CO). HRMS (ESI): Calculated for $C_{25}H_{34}N_2O_3Na$ [M+Na]$^+$: 433.2467; found: 433.2450.

***N*$^2$-(2-Cyanophenylcarbonyl)-*N*$^2$-octyl-*N*$^1$-phenyl-β-alaninamide (5).** Obtained from amine **26** (100 mg, 0.36 mmol), 2-cyanobenzoic acid (63 mg, 0.43 mmol), EDC (67 mg, 0.43 mmol) and HOBt (58 mg, 0.43 mmol) in 65% yield (96 mg). Chromatography: hexane/EtOAc, 1:1.

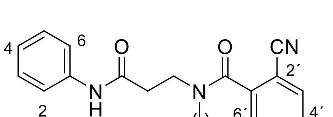


*R*$_f$ (hexane/EtOAc, 1:1): 0.50. IR (ATR, ν): 3308 (NH), 2229 (CN), 1729, 1687, 1546 (CON), 1601, 1498, 1442 (Ar). $^1$H-NMR (CDCl$_3$, δ): amide rotamers A:B 85:15; 0.85 (t, *J* = 7.0 Hz, 3H, CH$_3$), 1.08-1.29 (m, 10H, (C*H*$_2$)$_5$CH$_3$), 1.52 (m, 2H, C*H*$_2$(CH$_2$)$_5$CH$_3$ rotamer A), 1.60-1.73 (m, 2H, C*H*$_2$(CH$_2$)$_5$CH$_3$ rotamer B), 2.56 (t, *J* = 7.0 Hz, 2H, CH$_2$CO rotamer B), 2.83 (t, *J* = 6.4 Hz, 2H, CH$_2$CO rotamer A), 3.15 (t, *J* = 7.6 Hz, 2H, (CH$_2$)$_6$C*H*$_2$N rotamer A), 3.47-3.62 (m, 4H, (CH$_2$)$_6$C*H*$_2$N, COCH$_2$C*H*$_2$N rotamer B), 3.94 (t, *J* = 6.3 Hz, 2H, COCH$_2$C*H*$_2$N rotamer A), 7.08 (t, *J* = 7.4 Hz, 1H, $H_4$), 7.30 (t, *J* = 7.9 Hz, 2H, $H_3$, $H_5$), 7.43 (d, *J* = 7.6 Hz, 1H, $H_{6'}$), 7.50 (td, *J* = 7.7, 1.1 Hz, 1H, $H_{4'}$), 7.56 (d, *J* = 7.8 Hz, 2H, $H_2$, $H_6$), 7.64 (td, *J* = 7.7, 1.2 Hz, 1H, $H_{5'}$), 7.67 (d, *J* = 7.8 Hz, 1H, $H_{3'}$), 8.50 (br s, 1H, NH). $^{13}$C-NMR (CDCl$_3$, δ): 14.2 (CH$_3$), 22.6, 26.5, 28.8, 29.0 (2C), 31.8 ((*C*H$_2$)$_6$CH$_3$), 36.3 (*C*H$_2$CO), 42.3 (CH$_2$N), 50.1 ((CH$_2$)$_6$*C*H$_2$N), 109.9 ($C_{2'}$), 117.1 (CN), 120.0 ($C_2$, $C_6$), 124.2 ($C_4$), 126.9 ($C_{6'}$), 128.9 ($C_3$, $C_5$), 129.5 ($C_{5'}$), 132.8 ($C_{3'}$), 133.2 ($C_{4'}$), 138.3 ($C_1$), 140.7 ($C_{1'}$), 168.3, 169.8 (2CO). HRMS (ESI): Calculated for $C_{25}H_{30}N_3O_2$ [(M-H)$^-$]: 404.23435; found: 404.23578.

***N*$^2$-(3-Cyanophenylcarbonyl)-*N*$^2$-octyl-*N*$^1$-phenyl-β-alaninamide (6).** Obtained from amine **26** (50 mg, 0.18 mmol), 3-cyanobenzoic acid (32 mg, 0.21 mmol), EDC (10 mg, 0.21 mmol) and HOBt (12 mg, 0.21 mmol) in 66% yield (51 mg). Chromatography: hexane/EtOAc, 1:1.

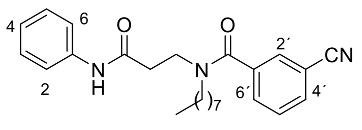


$R_f$ (hexane/EtOAc, 1:1): 0.35. IR (ATR, ν): 3308 (NH), 2232 (CN), 1687, 1619, 1545 (CON), 1601, 1579, 1499, 1442 (Ar). $^1$H-NMR ($CDCl_3$, δ): amide rotamers A:B 9:1; 0.89 (t, $J$ = 6.9 Hz, 3H, $CH_3$), 1.14-1.29 (m, 10H, ($CH_2$)$_5$$CH_3$), 1.54 (m, 2H, $CH_2$($CH_2$)$_5$$CH_3$ rotamer A), 1.69 (m, 2H, $CH_2$($CH_2$)$_5$$CH_3$ rotamer B), 2.54 (m, 2H, $CH_2$CO rotamer B), 2.83 (t, $J$ = 6.4 Hz, 2H, $CH_2$CO rotamer A), 3.26 (t, $J$ = 7.5 Hz, 2H, ($CH_2$)$_6$$CH_2$N rotamer A), 3.53 (m, 2H, ($CH_2$)$_6$$CH_2$N rotamer B), 3.65 (m, 2H, CO$CH_2$$CH_2$N rotamer B), 3.87 (t, $J$ = 6.4 Hz, 2H, CO$CH_2$$CH_2$N rotamer A), 7.13 (t, $J$ = 7.4 Hz, 1H, $H_4$), 7.32 (t, $J$ = 7.5 Hz, 2H, $H_3$, $H_5$), 7.50-7.60 (m, 4H, $H_2$, $H_6$, $H_{5'}$, $H_{6'}$), 7.65 (s, 1H, $H_{2'}$), 7.71 (d, $J$ = 7.4 Hz, 1H, $H_{4'}$), 8.76 (br s, 1H, NH). $^{13}$C-NMR ($CDCl_3$, δ): 14.2 ($CH_3$), 22.7, 26.5, 28.9, 29.1, 31.8 (($CH_2$)$_6$$CH_3$), 36.1 ($CH_2$CO), 42.8 ($CH_2$N), 50.7 (($CH_2$)$_6$$CH_2$N), 113.1 ($C_{3'}$), 117.9 (CN), 119.9 ($C_2$, $C_6$), 124.4 ($C_4$), 129.0 ($C_3$, $C_5$), 129.7 ($C_{5'}$/$C_{6'}$), 130.2 ($C_{2'}$), 130.8 ($C_{5'}$/$C_{6'}$), 133.1 ($C_{4'}$), 137.8, 138.1 ($C_1$, $C_{1'}$), 169.3, 170.2 (2CO). HRMS (ESI): Calculated for $C_{25}H_{31}N_3O_2Na$ [$(M+Na)^+$]: 428.23085; found: 428.23095.

***N*$^2$-(4-Cyanophenylcarbonyl)-*N*$^2$-octyl-*N*$^1$-phenyl-β-alaninamide (7).** Obtained from amine **26** (50 mg, 0.18 mmol), 4-cyanobenzoic acid (32 mg, 0.21 mmol), EDC (10 mg, 0.21 mmol) and HOBt (12 mg, 0.21 mmol) in 68% yield (55 mg). Chromatography: hexane/EtOAc, 1:1.

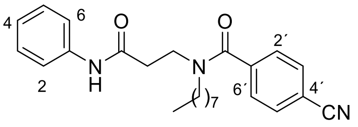


$R_f$ (hexane/EtOAc, 1:1): 0.30. Mp: 90-91 °C. IR (ATR, ν): 3311 (NH), 2232 (CN), 1686, 1611, 1545 (CON), 1498, 1439 (Ar). $^1$H-NMR ($CDCl_3$, δ): amide rotamers A:B, 9:1; 0.87 (t, $J$ = 6.9 Hz, 3H, $CH_3$), 1.10-1.27 (m, 10H, ($CH_2$)$_5$$CH_3$), 1.50 (m, 2H, $CH_2$($CH_2$)$_5$$CH_3$ rotamer A), 1.66 (m, 2H, $CH_2$($CH_2$)$_5$$CH_3$ rotamer B), 2.49 (m, 2H, $CH_2$CO rotamer B), 2.78 (t, $J$ = 6.4 Hz, 2H, $CH_2$CO rotamer A), 3.22 (t, $J$ = 7.8 Hz, 2H, ($CH_2$)$_6$$CH_2$N rotamer A), 3.50 (m, 2H, ($CH_2$)$_6$$CH_2$N rotamer B), 3.60 (m, 2H, CO$CH_2$$CH_2$N rotamer B), 3.84 (t, $J$ = 6.4 Hz, 2H, CO$CH_2$$CH_2$N rotamer A), 7.01 (t, $J$ = 7.3 Hz, 1H, $H_4$), 7.28 (t, $J$ = 7.8 Hz, 2H, $H_3$, $H_5$), 7.42 (d, $J$ = 8.1 Hz, 2H, $H_{3'}$, $H_{5'}$), 7.49 (d, $J$ = 8.0 Hz, 2H, $H_2$, $H_6$), 7.65 (d, $J$ = 8.1 Hz, 2H, $H_{2'}$, $H_{6'}$), 8.13 (br s, 1H, NH rotamer B), 9.02 (br s, 1H, NH rotamer A). $^{13}$C-NMR ($CDCl_3$, δ): 14.1 ($CH_3$), 22.6, 26.4, 28.8, 29.0 (2C), 31.7 (($CH_2$)$_6$$CH_3$), 35.8 ($CH_2$CO), 42.6 ($CH_2$N), 50.6 (($CH_2$)$_6$$CH_2$N), 113.4 ($C_{4'}$), 118.0 (CN), 119.9 ($C_2$, $C_6$), 124.2 ($C_4$), 127.2 ($C_{3'}$, $C_{5'}$), 128.9 ($C_3$, $C_5$), 132.5 ($C_{2'}$, $C_{6'}$), 138.2 ($C_1$), 140.7 ($C_{1'}$), 169.3, 170.6 (2CO). HRMS (ESI): Calculated for $C_{25}H_{31}N_3O_2Na$ [$(M+Na)^+$]: 428.23085; found: 428.23099. Elemental analysis calculated for $C_{25}H_{31}N_3O_2$: C, 74.04; H, 7.70; N, 10.36; found: C, 73.81; H, 7.54; N, 10.19.

***N*$^2$-(4-Fluorophenylcarbonyl)-*N*$^2$-octyl-*N*$^1$-phenyl-β-alaninamide (8).** Obtained from amine **26** (100 mg, 0.36 mmol), 4-fluorobenzoic acid (59 mg, 0.42 mmol), EDC (67 mg, 0.43 mmol) and HOBt (58 mg, 0.43 mmol) in 75% yield (108 mg). Chromatography: hexane/EtOAc, 7:3 to 3:7.

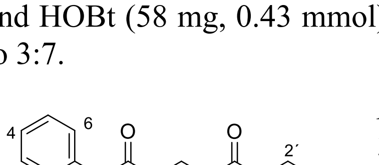


$R_f$ (hexane/EtOAc, 1:1): 0.42. Mp: 76-77 °C. IR (ATR, ν): 3306, 3136 (NH), 1688, 1666, 1546 (CON), 1602, 1499, 1442 (Ar). $^1$H-NMR ($CDCl_3$, δ): 0.86 (t, $J$ = 6.9 Hz, 3H, $CH_3$), 1.11-1.26 (m, 10H, ($CH_2$)$_5$$CH_3$), 1.51 (m, 2H, $CH_2$($CH_2$)$_5$$CH_3$), 2.75 (t, $J$ = 5.9 Hz, 2H, $CH_2$CO), 3.26 (t, $J$ = 7.2 Hz, 2H, ($CH_2$)$_6$$CH_2$N), 3.82 (t, $J$ = 6.0 Hz, 2H, CO$CH_2$$CH_2$N), 7.03 (t, $J$ = 8.6 Hz, 2H, $H_{2'}$, $H_{6'}$), 7.07 (t, $J$ = 7.4 Hz, 1H, $H_4$), 7.27 (t, $J$ = 7.9 Hz, 2H, $H_3$, $H_5$), 7.32 (dd, $J$ = 8.4, 5.5 Hz, 2H, $H_{3'}$, $H_{5'}$), 7.52 (d, $J$ = 7.9 Hz, 2H, $H_2$, $H_6$), 9.29 (br s, 1H, NH). $^{13}$C-NMR ($CDCl_3$, δ): 14.2 ($CH_3$), 22.7, 26.5, 28.9, 29.1 (2C), 31.8 (($CH_2$)$_6$$CH_3$), 36.1 ($CH_2$CO), 42.8 ($CH_2$N), 50.8 (($CH_2$)$_6$$CH_2$N), 115.7 ($C_{3'}$, $C_{5'}$), 119.9 ($C_2$, $C_6$), 124.1 ($C_4$), 128.7 (d, $J$ = 8.4 Hz, $C_{2'}$, $C_{6'}$), 128.9 ($C_3$, $C_5$), 132.5 ($C_{1'}$), 138.2 ($C_1$), 163.3

($C_{4'}$), 169.7, 171.8 (2CO). MS (ESI): 399.1 [$(M+H)^+$]. Elemental analysis calculated for $C_{26}H_{36}FN_3O_2$: C, 72.33; H, 7.84; N, 7.03; found: C, 72.41; H, 7.78; N, 6.98.

***N*²-[4-(Trifluoromethyl)phenylcarbonyl]-*N*²-octyl-*N*¹-phenyl-β-alaninamide (9).** Obtained from amine **26** (100 mg, 0.36 mmol), 4-(trifluoromethyl)benzoic acid (82 mg, 0.43 mmol), EDC (66 mg, 0.43 mmol) and HOBt (49 mg, 0.43 mmol) in 78% yield (126 mg). Chromatography: hexane/EtOAc, 7:3 to 1:1.

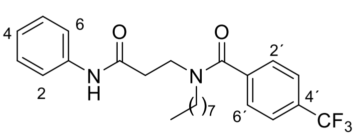

$R_f$ (hexane/EtOAc, 1:1): 0.53. Mp: 80-81 ºC. IR (ATR, ν): 3307 (NH), 1688, 1666, 1615, 1546 (CON), 1499, 1443 (Ar). $^1$H-NMR ($CDCl_3$, δ): amide rotamers A:B, 9:1; 0.88 (t, $J$ = 6.9 Hz, 3H, $CH_3$), 1.12-1.28 (m, 10H, $(CH_2)_5CH_3$), 1.53 (m, 2H, $CH_2(CH_2)_5CH_3$ rotamer A), 1.69 (m, 2H, $CH_2(CH_2)_5CH_3$ rotamer B), 2.51 (m, 2H, $CH_2CO$ rotamer B), 2.83 (t, $J$ = 6.4 Hz, 2H, $CH_2CO$ rotamer A), 3.26 (t, $J$ = 7.5 Hz, 2H, $(CH_2)_6CH_2N$ rotamer A), 3.53 (m, 2H, $(CH_2)_6CH_2N$ rotamer B), 3.64 (m, 2H, $COCH_2CH_2N$ rotamer B), 3.88 (t, $J$ = 6.4 Hz, 2H, $COCH_2CH_2N$ rotamer A), 7.10 (t, $J$ = 7.4 Hz, 1H, $H_4$), 7.28 (t, $J$ = 7.8 Hz, 2H, $H_3$, $H_5$), 7.45 (d, $J$ = 8.0 Hz, 2H, $H_{2'}$, $H_{6'}$ rotamer A), 7.50 (d, $J$ = 8.0 Hz, 2H, $H_2$, $H_6$), 7.64 (d, $J$ = 8.1 Hz, 2H, $H_{3'}$, $H_{5'}$ rotamer A), 7.72 (d, $J$ = 8.2 Hz, 2H, $H_{2'}$, $H_{6'}$ rotamer B), 8.17 (d, $J$ = 8.1 Hz, 2H, $H_{3'}$, $H_{5'}$ rotamer B), 9.25 (br s, 1H, NH). $^{13}$C-NMR ($CDCl_3$, δ): 14.1 ($CH_3$), 22.6, 26.4, 28.8, 29.0, 29.1, 31.7 ($(CH_2)_6CH_3$), 35.9 ($CH_2CO$), 42.8 ($CH_2N$), 50.7 ($(CH_2)_6CH_2N$), 120.0 ($C_2$, $C_6$), 124.2 ($C_4$), 125.7 (q, $J$ = 3.6 Hz, $C_{3'}$, $C_{5'}$), 126.9 ($C_{2'}$, $C_{6'}$), 127.3 ($CF_3$), 128.9 ($C_3$, $C_5$), 131.7 ($C_{4'}$), 138.3 ($C_1$), 139.9 ($C_{1'}$), 169.6, 171.3 (2CO). MS (ESI): 449.2 [$(M+H)^+$]. Elemental analysis calculated for $C_{25}H_{31}F_3N_2O_2$: C, 66.95; H, 6.97; N, 6.25; found: C, 66.23; H, 6.73; N, 6.01.

***N*²-(4-Acetamidophenylcarbonyl)-*N*²-octyl-*N*¹-phenyl-β-alaninamide (10).** Obtained from amine **26** (85 mg, 0.31 mmol), 4-acetamidobenzoic acid (66 mg, 0.37 mmol), EDC (57 mg, 0.37 mmol) and HOBt (50 mg, 0.37 mmol) in 80% yield (108 mg). Chromatography: hexane/EtOAc, 2:8.

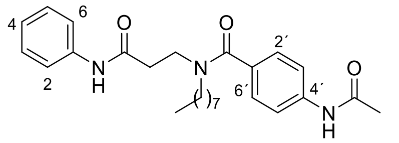

$R_f$ (hexane/EtOAc, 2:8): 0.19. IR (ATR, ν): 3300 (NH), 1669, 1535 (CON), 1602, 1468, 1438 (Ar). $^1$H-NMR ($CDCl_3$, δ): amide rotamers A:B, 8:2; 0.85 (t, $J$ = 6.8 Hz, 3H, $CH_3CH_2$), 1.16-1.29 (m, 10H, $(CH_2)_5CH_3$), 1.49-1.62 (m, 2H, $CH_2(CH_2)_5CH_3$ rotamers A and B), 1.97 (s, 3H, $CH_3CO$ rotamer B), 2.15 (s, 3H, $CH_3CO$ rotamer A), 2.22 (m, 2H, $CH_2CO$ rotamer B), 2.76 (m, 2H, $CH_2CO$ rotamer A), 3.26 (m, 2H, $(CH_2)_6CH_2N$ rotamer A), 3.45 (m, 2H, $(CH_2)_6CH_2N$ rotamer B), 3.60 (m, 2H, $COCH_2CH_2N$ rotamer B), 3.83 (m, 2H, $COCH_2CH_2N$ rotamer A), 7.07 (t, $J$ = 7.4 Hz, 1H, $H_4$), 7.20 (d, $J$ = 7.6 Hz, 2H, $H_{2'}$, $H_{6'}$), 7.27 (t, $J$ = 7.8 Hz, 2H, $H_3$, $H_5$), 7.44 (d, $J$ = 7.4 Hz, 2H, $H_{3'}$, $H_{5'}$), 7.53 (d, $J$ = 7.7 Hz, 2H, $H_2$, $H_6$), 8.15 (br s, 1H, NH), 9.19 (br s, 1H, NH). $^{13}$C-NMR ($CDCl_3$, δ): 14.2 ($CH_3CH_2$), 22.7 ($CH_2$), 24.6 ($CH_3CO$), 26.6, 26.7, 29.0, 29.2, 31.8 (5$CH_2$), 36.3 ($CH_2CO$), 42.6 ($CH_2N$), 50.8 ($(CH_2)_6CH_2N$), 119.7 ($C_{3'}$, $C_{5'}$), 120.1 ($C_2$, $C_6$), 124.2 ($C_4$), 127.6 ($C_{2'}$, $C_{6'}$), 129.0 ($C_3$, $C_5$), 131.8 ($C_{1'}$), 138.4 ($C_1$), 139.5 ($C_{4'}$), 169.0, 169.1, 169.9 (3CO). HRMS (ESI): Calculated for $C_{26}H_{35}N_3O_3Na$ [$(M+Na)^+$]: 460.2570; found: 460.25833.

***N*²-Octyl-*N*¹-phenyl-*N*²-(tetrahydrofuran-3-ylcarbonyl)-β-alaninamide (11).** Obtained from amine **26** (100 mg, 0.36 mmol), tetrahydrofuran-3-carboxylic acid (84 mg, 0.72 mmol), EDC (138 mg, 0.72 mmol) and HOBt (97 mg, 0.72 mmol) in 84% yield (113 mg). Chromatography: hexane/EtOAc, 7:3.

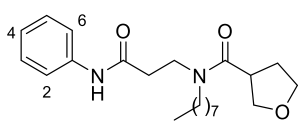

$R_f$ (hexane/EtOAc, 7:3): 0.13. Mp: 68-70 ºC. IR (ATR, ν): 3309 (NH), 1684, 1545, (CON), 1624, 1494, 1442 (Ar). $^1$H-NMR

($CDCl_3$, δ): 0.89 (t, $J$ = 6.7 Hz, 3H, $CH_3$), 1.28-1.30 (m, 10H, $(CH_2)_5CH_3$), 1.56-1.61 (m, 2H, $CH_2(CH_2)_5CH_3$), 2.01-2.23 (m, 2H, $CH_2CH_2O$), 2.69 (t, $J$ = 6.6 Hz, 2H, $CH_2CO$), 3.22 (qt, $J$ = 7.6 Hz, 1H, CH), 3.34 (t, $J$ = 6.8 Hz, 2H, $NCH_2(CH_2)_6$), 3.70 (t, $J$ = 6.6 Hz, 2H, $NCH_2CH_2CO$), 3.81-4.02 (m, 4H, $2CH_2O$), 7.09 (t, $J$ = 7.4 Hz, 1H, $H_4$), 7.31 (t, $J$ = 7.9 Hz, 2H, $H_3$,$H_5$), 7.54 (d, $J$ = 7.9 Hz, 2H, $H_2$, $H_6$), 8.47 (br s, 1H, NH). $^{13}$C-NMR ($CDCl_3$, δ): 14.0 ($CH_3$), 22.6, 26.8, 29.2, 29.3, 29.5 ($(CH_2)_5CH_3$), 30.8 ($CH_2CH_2O$), 31.7 ($CH_2(CH_2)_5$), 36.8 ($CH_2CO$), 41.2 (CH), 43.2 ($NCH_2CH_2CO$), 49.0 ($NCH_2(CH_2)_6$), 68.6, 71.2 ($2CH_2O$), 119.8 ($C_2$, $C_6$), 124.1 ($C_4$), 128.9 ($C_3$, $C_5$), 138.2 ($C_1$), 168.4, 174.1 (2CO). MS (ESI): 375.2 [$(M+H)^+$]. Elemental analysis calculated for $C_{22}H_{34}N_2O_3$: C, 70.55; H, 9.15; N, 7.48; found: C, 70.83; H, 8.77; N, 7.50.

***N*²-(2-Furoyl)-*N*²-octyl-*N*¹-phenyl-β-alaninamide (12).** Obtained from amine **26** (85 mg, 0.31 mmol), 2-furoic acid (76 mg, 0.62 mmol), EDC (96 mg, 0.62 mmol) and HOBt (84 mg, 0.62 mmol) in 95% yield (109 mg). Chromatography: hexane/EtOAc, 1:1.

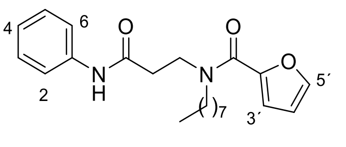


$R_f$ (hexane/EtOAc, 1:1): 0.40. Mp: 91-93 °C. IR (ATR, ν): 3315 (NH), 1668 (CON), 1605 (Ar). $^1$H-NMR ($CDCl_3$, δ): 0.87 (t, $J$ = 6.6 Hz, 3H, $CH_3$), 1.25 (m, 10H, $(CH_2)_5CH_3$), 1.65 (m, 2H, $CH_2(CH_2)_5CH_3$), 2.75 (t, $J$ = 6.3 Hz, 2H, $CH_2CO$), 3.61 (m, 2H, $(CH_2)_6CH_2N$), 3.83 (m, 2H, $COCH_2CH_2N$), 6.43 (m, 1H, $H_{4'}$), 6.95 (m, 1H, $H_{3'}$), 7.06 (t, $J$ = 7.4 Hz, 1H, $H_4$), 7.27 (t, $J$ = 7.8 Hz, 2H, $H_3$, $H_5$), 7.41 (m, 1H, $H_{5'}$), 7.49-7.60 (m, 2H, $H_2$, $H_6$), 9.29 (br s, 1H, NH). $^{13}$C-NMR ($CDCl_3$, δ): 14.1 ($CH_3$), 22.6, 26.8, 29.2, 29.3, 29.7, 31.8 ($(CH_2)_6CH_3$), 36.2 ($CH_2CO$), 44.3 ($CH_2N$), 49.6 ($(CH_2)_6CH_2N$), 111.4, 116.5 ($C_{3'}$, $C_{4'}$), 119.9 ($C_2$, $C_6$), 123.9 ($C_4$), 128.8 ($C_3$, $C_5$), 138.5 ($C_1$), 144.0 ($C_{5'}$), 147.9 ($C_{2'}$), 160.5 (CON), 169.6 (CONH). MS (ESI): 371.2 [$(M+H)^+$]. Elemental analysis calculated for $C_{22}H_{30}N_2O_3$: C, 71.32; H, 8.16; N, 7.56; found: C, 71.35; H, 8.03; N, 7.30.

***N*²-(3-Furoyl)-*N*²-octyl-*N*¹-phenyl-β-alaninamide (13).** Obtained from amine **26** (100 mg, 0.36 mmol), 3-furoic acid (81 mg, 0.72 mmol), EDC (138 mg, 0.72 mmol) and HOBt (97 mg, 0.72 mmol) in 98% yield (128 mg). Chromatography: hexane/EtOAc, 7:3.

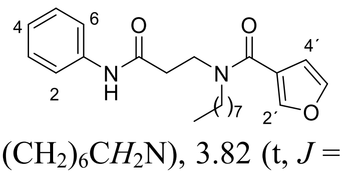


$R_f$ (hexane/EtOAc, 7:3): 0.14. IR (ATR, ν): 3281 (NH), 1685 (CON), 1605, 1547, 1501, 1441 (Ar). $^1$H-NMR ($CDCl_3$, δ): 0.88 (t, $J$ = 6.7 Hz, 3H, $CH_3$), 1.25 (m, 10H, $(CH_2)_5CH_3$), 1.61 (m, 2H, $CH_2(CH_2)_5CH_3$), 2.76 (m, 2H, $CH_2CO$), 3.44 (t, $J$ = 7.9 Hz, 2H, $(CH_2)_6CH_2N$), 3.82 (t, $J$ = 6.6 Hz, 2H, $COCH_2CH_2N$), 6.55 (m, 1H, $H_{4'}$), 7.08 (t, $J$ = 7.4 Hz, 1H, $H_4$), 7.29 (t, $J$ = 8.3 Hz, 2H, $H_3$, $H_5$), 7.41 (t, $J$ = 1.7 Hz, 1H, $H_{5'}$), 7.55 (d, $J$ = 7.8 Hz, 2H, $H_2$, $H_6$), 7.68 (m, 1H, $H_{2'}$). $^{13}$C-NMR ($CDCl_3$, δ): 14.1 ($CH_3$), 22.6, 26.6, 29.2 (3C), 31.7 ($(CH_2)_6CH_3$), 36.5 ($CH_2CO$), 43.2 ($CH_2N$), 50.1 ($(CH_2)_6CH_2N$), 110.1 ($C_{4'}$), 119.8 ($C_2$, $C_6$), 121.3 ($C_{3'}$), 124.1 ($C_4$), 128.9 ($C_3$, $C_5$), 138.3 ($C_1$), 143.1, 143.2 ($C_{2'}$, $C_{5'}$), 165.6 (CON), 169.4 (CONH). MS (ESI): 371.2 [$(M+H)^+$]. Elemental analysis calculated for $C_{22}H_{30}N_2O_3$: C, 71.32; H, 8.16; N, 7.56; found: C, 71.15; H, 7.91; N, 7.61.

***N*²-Octyl-*N*¹-phenyl-*N*²-(1*H*-pyrrol-2-ylcarbonyl)-β-alaninamide (15).** Obtained from amine **26** (85 mg, 0.31 mmol), 1*H*-2-pyrrolcarboxylic acid (70 mg, 0.62 mmol), EDC (95 mg, 0.62 mmol) and HOBt (84 mg, 0.62 mmol) in 51% yield (58 mg). In this case the carboxylic acid, the coupling reagents and the amine **26** were all dissolved in DCM at the same time to avoid the acylation of the pyrrole ring. Chromatography: hexane/EtOAc, 7:3.

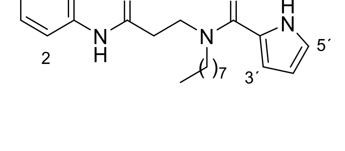


$R_f$ (hexane/EtOAc, 1:1): 0.40. Mp: 105-107 °C. IR (ATR, ν): 3264 (NH), 1666 (CON), 1599, 1548, 183, 1445 (Ar). $^1$H-NMR ($CDCl_3$, δ): 0.88 (t, $J$ = 6.7 Hz, 3H, $CH_3$), 1.28-1.31 (m, 10H, $(CH_2)_5CH_3$), 1.71 (m, 2H, $CH_2(CH_2)_5CH_3$), 2.74 (t, $J$ = 6.7 Hz,

2H, $CH_2CO$), 3.58 (m, 2H, $(CH_2)_6CH_2N$), 3.87 (m, 2H, $COCH_2CH_2N$), 6.25-6.28 (m, 1H, $H_{4'}$), 6.54 (m, 1H, $H_{3'}$), 6.90 (m, 1H, $H_{5'}$), 7.07 (t, $J$ = 7.3 Hz, 1H, $H_4$), 7.28 (t, $J$ = 7.8 Hz, 2H, $H_3$, $H_5$), 7.55 (d, $J$ = 7.8 Hz, 2H, $H_2$, $H_6$), 8.84 (br s, 1H, NH), 9.84 (br s, 1H, NH). $^{13}$C-NMR ($CDCl_3$, δ): 14.1 ($CH_3$), 22.6, 26.8, 28.6, 29.2, 29.4, 31.8 ($(CH_2)_6CH_3$), 36.7 ($CH_2CO$), 44.3 ($CH_2N$), 49.3 ($(CH_2)_6CH_2N$), 110.3, 112.2 ($C_{3'}$, $C_{4'}$), 119.9 ($C_2$, $C_6$), 121.3 ($C_{5'}$), 124.1 ($C_4$), 124.4 ($C_{2'}$), 128.9 ($C_3$, $C_5$), 138.3 ($C_1$), 162.5 (CON), 169.4 (CONH). MS (ESI): 370.2 $[M+H]^+$. Elemental analysis calculated for $C_{22}H_{30}N_2O_3$: C, 71.51; H, 8.46; N, 11.37; found: C, 71.29; H, 8.11; N, 11.35.

***N*$^2$-Octyl-*N*$^1$-phenyl-*N*$^2$-(1*H*-pyrrol-3-ylcarbonyl)-β-alaninamide (16).** Obtained from amine **26** (80 mg, 0.29 mmol), 1*H*-3-pyrrolcarboxilic acid (65 mg, 0.58 mmol), EDC (90 mg, 0.58 mmol) and HOBt (78 mg, 0.58 mmol) in 54% yield (58 mg). In this case the carboxylic acid, the coupling reagents and the amine **26** were all dissolved in DCM at the same time to avoid the acylation of the pyrrole ring. Chromatography: hexane/EtOAc, 3:7.

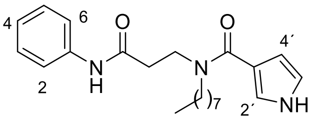


$R_f$ (EtOAc): 0.25. Mp: 99-101 °C. IR (ATR, ν): 3250 (NH), 1598 (CON), 1548 (Ar). $^1$H-NMR ($CDCl_3$, δ): 0.79 (t, $J$ = 6.7 Hz, 3H, $CH_3$), 1.16 (m, 10H, $(CH_2)_5CH_3$), 1.54 (m, 2H, $CH_2(CH_2)_5CH_3$), 2.61 (m, 2H, $CH_2CO$), 3.41 (t, $J$ = 7.2 Hz, 2H, $(CH_2)_6CH_2N$), 3.74 (t, $J$ = 6.6 Hz, 2H, $COCH_2CH_2N$), 6.26 (m, 1H, $H_{4'}$), 6.56 (m, 1H, $H_{5'}$), 6.92 (m, 1H, $H_{2'}$), 6.97 (t, $J$ = 7.4 Hz, 1H, $H_4$), 7.18 (t, $J$ = 7.8 Hz, 2H, $H_3$, $H_5$), 7.49 (d, $J$ = 7.9 Hz, 2H, $H_2$, $H_6$), 9.37 (br s, 2H, 2NH). $^{13}$C-NMR ($CDCl_3$, δ): 14.1 ($CH_3$), 22.6, 26.7, 28.9, 29.2, 29.3, 31.8 ($(CH_2)_6CH_3$), 36.6 ($CH_2CO$), 43.4 ($CH_2N$), 49.8 ($(CH_2)_6CH_2N$), 108.6 ($C_{4'}$), 118.3 ($C_{5'}$), 118.4 ($C_{3'}$), 120.0 ($C_2$, $C_6$), 121.1 ($C_{2'}$), 123.9 ($C_4$), 128.8 ($C_3$, $C_5$), 138.5 ($C_1$), 168.4 (CON), 170.0 (CONH). MS (ESI): 370.2 $[M+H]^+$. Elemental analysis calculated for $C_{22}H_{30}N_2O_3$: C, 71.51; H, 8.46; N, 11.37; found: C, 71.40; H, 8.22; N, 11.18.

***N*$^2$-[(1-Methyl-1*H*-pyrrol-2-yl)carbonyl]-*N*$^2$-octyl-*N*$^1$-phenyl-β-alaninamide (17).** Obtained from amine **26** (163 mg, 0.59 mmol), 1-methyl-1*H*-pyrrol-2-carboxylic acid (150 mg, 1.2 mmol), EDC (230 mg, 1.2 mmol) and HOBt (162 mg, 1.2 mmol) in 49% yield (111 mg). Chromatography: hexane/EtOAc, 7:3.

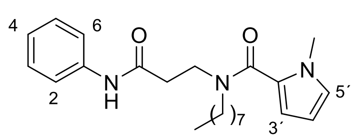


$R_f$ (hexane/EtOAc, 7:3): 0.44. IR (ATR, ν): 3277 (NH), 1684 (CO), 1602, 1543, 1473, 1439 (Ar). $^1$H-NMR ($CDCl_3$, δ): 0.87 (t, $J$ = 6.7 Hz, 3H, $CH_3CH_2$), 1.23-1.30 (m, 10H, $(CH_2)_5CH_3$), 1.58-1.65 (m, 2H, $CH_2(CH_2)_5CH_3$), 2.73 (t, $J$ = 6.8 Hz, 2H, $CH_2CO$), 3.54 (t, $J$ = 7.6 Hz, 2H, $NCH_2(CH_2)_6$), 3.67 (s, 3H, $CH_3N$), 3.80 (t, $J$ = 6.8 Hz, 2H, $NCH_2CH_2CO$), 6.06 (dd, $J$ = 3.7, 2.7 Hz, 1H, $H_{4'}$), 6.30 (dd, $J$ = 3.7, 1.4 Hz, 1H, $H_{3'}$), 6.66 (t, $J$ = 2.9 Hz, 1H, $H_{5'}$), 7.07 (t, $J$ = 7.4 Hz, 1H, $H_4$), 7.28 (t, $J$ = 7.8 Hz, 2H, $H_3$, $H_5$), 7.55 (d, $J$ = 8.1 Hz, 2H, $H_2$, $H_6$), 9.11 (br s, 1H, NH). $^{13}$C-NMR ($CDCl_3$, δ): 14.5 ($CH_3CH_2$), 23.0, 26.9, 29.2, 29.6 (2C), 32.1 ($(CH_2)_6CH_3$), 35.9 ($CH_3N$), 36.8 ($CH_2CO$), 43.5 ($NCH_2CH_2CO$), 50.6 ($NCH_2(CH_2)_6$), 107.5, 112.5 ($C_{3'}$, $C_{4'}$), 120.2 ($C_2$, $C_6$), 124.4 ($C_4$), 125.7 ($C_{2'}$), 126.6 ($C_{5'}$), 129.2 ($C_3$, $C_5$), 138.8 ($C_1$), 165.6 (CON), 170.1 (CONH). HRMS (ESI, m/z): calculated for $C_{23}H_{34}N_3O_2$ $[M+H]^+$: 384.2646; found: 384.2655.

***N*$^2$-(1*H*-Imidazol-2-ylcarbonyl)-*N*$^2$-octyl-*N*$^1$-phenyl-β-alaninamide (18).** Obtained from amine **26** (95 mg, 0.33 mmol), 1*H*-imidazol-2-carboxylic acid (150 mg, 0.66 mmol), EDC (118 mg, 0.66 mmol) and HOBt (64 mg, 0.66 mmol) in 60% yield (71 mg). Chromatography: hexane/EtOAc, 7:3.

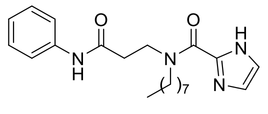


$R_f$ (hexane/EtOAc, 7:3): 0.44. Mp: 96-97 ºC. IR (ATR, ν): 3201 (NH), 1668 (CO), 1604, 1546, 1483, 1443 (Ar). $^1$H-NMR ($CDCl_3$, δ): amide romaters A:B 6:1, 0.81 (t, $J$ = 6.7 Hz, 3H, $CH_3$), 1.20-1.26 (m, 10H,

$(CH_2)_5CH_3$), 1.60 (m, 2H, $CH_2(CH_2)_5CH_3$), 2.71 (m, 2H, $CH_2CO$, rotamer B), 2.82 (t, *J* = 8.1 Hz, 2H, $CH_2CO$, rotamer A), 3.47 (t, *J* = 7.5 Hz, 2H, $NCH_2(CH_2)_6$, rotamer A), 3.81 (m, 2H, $NCH_2CH_2CO$, rotamer B), 4.21 (m, 2H, $NCH_2(CH_2)_6$, rotamer B), 4.30 (t, *J* = 8.0 Hz, 2H, $NCH_2CH_2CO$, rotamer A), 7.03 (t, *J* = 7.4 Hz, 1H, $H_4$), 7.16 (m, 1H, $CH_{HetAr}$), 7.19 (s, 1H, $CH_{HetAr}$), 7.25-7.30 (m, 2H, $H_3$, $H_5$), 7.45-7.55 (m, 2H, $H_2$, $H_6$), 8.29 (br s, 1H, NH, rotamer B), 9.98 (br s, 1H, NH, rotamer A), 10.94 (br s, 1H, NH, rotamer B), 11.52 (br s, 1H, NH, rotamer A). $^{13}$C-NMR ($CDCl_3$, δ): 14.1 ($CH_3$), 22.6, 27.1, 27.8, 29.2, 29.4, 31.8 ($(CH_2)_6CH_3$), 39.7 ($CH_2CO$), 46.8 ($NCH_2CH_2CO$), 48.3 ($NCH_2(CH_2)_6$), 118.8 ($CH_{HetAr}$), 119.6 ($C_2$, $C_6$), 123.9 ($C_4$), 129.1 ($C_3$, $C_5$), 129.8 ($CH_{HetAr}$), 138.6 ($C_1$), 141.5 ($C_{1'}$), 158.3 (CON), 169.1 (CONH). MS (ESI): 371.2 [$(M+H)^+$]. Elemental analysis calculated for $C_{21}H_{30}N_4O_2$ C, 68.08; H, 8.16; N, 15.12; found: C, 68.42; H, 8.49; N, 15.18.

***N*²-Octyl-*N*¹-phenyl-*N*²-(1*H*-1,2,4-triazol-3-ylcarbonyl)-β-alaninamide (19).** Obtained from amine **26** (150 mg, 0.54 mmol), 1*H*-1,2,4-triazol-3-carboxylic acid (124 mg, 1.1 mmol), EDC (171 mg, 1.1 mmol) and HOBt (149 mg, 1.1 mmol) in 45% yield (90 mg). Chromatography: EtOAc.

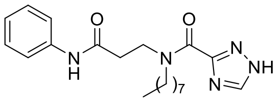

$R_f$ (EtOAc): 0.25. Mp: 110-111 ºC. IR (ATR, ν): 3281 (NH), 1623 (CO), 1547 (Ar). $^1$H-NMR ($CDCl_3$, δ): amide rotamers A:B 3:2, 0.84-0.89 (m, 3H, $CH_3$), 1.25-1.29 (m, 10H, $(CH_2)_5CH_3$), 1.61-1.78 (m, 2H, $CH_2(CH_2)_5CH_3$), 2.81-2.87 (m, 2H, $CH_2CO$), 3.55 (t, *J* = 7.6 Hz, 2H, $NCH_2(CH_2)_6$, rotamer B), 3.92 (t, *J* = 7.3 Hz, 2H, $NCH_2CH_2CO$, rotamer A), 4.08-4.15 (m, 2H, $NCH_2(CH_2)_6$, rotamer A), 4.27 (t, *J* = 7.7 Hz, 2H, $NCH_2CH_2CO$, rotamer B), 7.05-7.12 (m, 1H, $H_4$), 7.31 (t, *J* = 8.1 Hz, 2H, $H_3$, $H_5$), 7.54-7.58 (m, 2H, $H_2$, $H_6$), 8.13 (s, 1H, $H_{5'}$, rotamer A), 8.22 (m, 1H, $CH_{HetAr}$, rotamer B), 8.96 (br s, 1H, NH), 13.94 (br s, 1H, NH). $^{13}$C-NMR ($CDCl_3$, δ): 14.1 ($CH_3$), 22.6, 26.6, 27.0, 27.6, 29.2 (2C), 29.3, 31.8 ($(CH_2)_6CH_3$, rotamers A and B), 35.7, 38.9 ($\underline{C}H_2CO$, rotamers A and B), 45.6, 46.1 ($N\underline{C}H_2CH_2CO$, rotamers A and B), 48.1, 50.2 ($N\underline{C}H_2(CH_2)_6$, rotamers A and B), 119.7, 120.0 ($C_2$, $C_6$, rotamers A and B), 124.3 ($C_4$), 128.9, 129.0 ($C_3$, $C_5$, rotamers A and B), 138.1, 138.2 ($C_1$, rotamers A and B), 150.8 ($C_{5'}$), 158.8 ($C_{3'}$), 168.7, 169.0 (CO, rotamers A and B), 171.2 (CO). MS (ESI): 372.2 [$(M+H)^+$]. Elemental analysis calculated for $C_{20}H_{29}N_5O_2$ C, 64.66; H, 7.87; N, 18.85; found: C, 64.49; H, 7.66; N, 18.39.

***N*²-Octyl-*N*¹-phenyl-*N*²-(1,3-oxazol-5-ylcarbonyl)-β-alaninamide (20).** Obtained from amine **26** (120 mg, 0.43 mmol), 1,3-oxazol-5-carboxylic acid (97 mg, 0.86 mmol), EDC (133 mg, 0.86 mmol) and HOBt (116 mg, 0.86 mmol) in 95% yield (150 mg). Chromatography: hexane/EtOAc, 1:1.

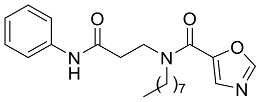

$R_f$ (hexane/EtOAc, 1:1): 0.21. IR (ATR, ν): 3312 (NH), 1684 (C=O), 1623, 1546, 1495, 1442 (Ar). $^1$H-NMR ($CDCl_3$, δ): 0.88 (t, *J* = 6.7 Hz, 3H, $CH_3$), 1.27 (m, 10H, $(CH_2)_5CH_3$), 1.66 (m, 2H, $CH_2(CH_2)_5CH_3$), 2.79 (m, 2H, $CH_2CO$), 3.59 (m, 2H, $NCH_2(CH_2)_6$), 3.85 (m, 2H, $NCH_2CH_2CO$), 7.08 (t, *J* = 7.3 Hz, 1H, $H_4$), 7.28 (t, *J* = 7.8 Hz, 2H, $H_3$, $H_5$), 7.54-7.56 (m, 3H, $H_2$, $H_6$, $CH_{4'}$), 7.91 (m, 1H, $CH_{2'}$), 8.52 y 8.94 (br s, 1H, NH). $^{13}$C-NMR ($CDCl_3$, δ): 13.0 ($CH_3$), 21.6, 25.6, 28.1, 28.2, 28.4, 30.7 ($(CH_2)_6CH_3$), 34.8 ($CH_2CO$), 43.3 ($NCH_2CH_2CO$), 48.7 ($NCH_2(CH_2)_6$), 118.9 ($C_2$, $C_6$), 123.2 ($C_4$), 127.9 ($C_3$, $C_4$), 130.4 ($C_{4'}$), 137.2 ($C_1$), 144.2 ($C_{5'}$), 150.7 ($C_{2'}$), 157.6 (CON), 168.3 (CONH). HRMS (ESI): Calculada para $C_{21}H_{29}N_3O_3Na$ [$(M+Na)^+$]: 394.2101. Encontrada: 394.2908.

***N*²-Octyl-*N*¹-phenyl-*N*²-(pyridin-3-ylcarbonyl)-β-alaninamide (22).** Obtained from amine **26** (100 mg, 0.36 mmol), nicotinic acid (87 mg, 0.72 mmol), EDC (138 mg, 0.72 mmol) and HOBt (97 mg, 0.72 mmol) in 36% yield (50 mg). Chromatography: hexane/EtOAc, 2:8.

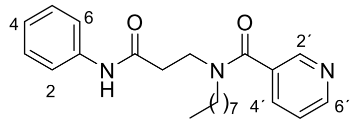

$R_f$ (EtOAc): 0.28. IR (ATR, ν): 3312 (NH), 1685 (CON), 1612, 1546, 1495, 1440 (Ar). $^1$H-NMR ($CDCl_3$, δ): amide rotamers A:B, 6:1; 0.87 (t, $J$ = 6.8 Hz, 3H, $CH_3$), 1.13-1.18 (m, 10H, $(CH_2)_5CH_3$), 1.53 (m, 2H, $CH_2(CH_2)_5CH_3$), 2.50 (m, 2H, $CH_2CO$, rotamer B), 2.79 (t, $J$ = 5.7 Hz, 2H, $CH_2CO$, rotamer A), 3.28 (t, $J$ = 7.0 Hz, 2H, $(CH_2)_6CH_2N$, rotamer A), 3.49 (m, 2H, $(CH_2)_6CH_2N$, rotamer B), 3.64 (m, 2H, $COCH_2CH_2N$, rotamer B), 3.86 (m, 2H, $COCH_2CH_2N$, rotamer A), 7.07 (t, $J$ = 7.3 Hz, 1H, $H_4$), 7.24-7.32 (m, 3H, $H_3$, $H_5$, $H_{5'}$), 7.53 (d, $J$ = 7.6 Hz, 2H, $H_2$, $H_6$), 7.67 (d, $J$ = 7.7 Hz, 1H, $H_{4'}$), 8.64 (d, $J$ = 1.4 Hz, 2H, $H_{2'}$, $H_{6'}$), 9.27 (br s, 1H, NH). $^{13}$C-NMR ($CDCl_3$, δ): 14.0 ($CH_3$), 22.5, 26.4, 28.9, 29.0 (2C), 31.6 ($(CH_2)_6CH_3$), 35.8 ($CH_2CO$), 42.7 ($CH_2N$), 50.7 ($(CH_2)_6CH_2N$), 119.9 ($C_2$, $C_6$), 123.4 ($C_{5'}$), 124.1 ($C_4$), 128.8 ($C_3$, $C_5$), 132.4 ($C_{3'}$), 134.2 ($C_{4'}$), 138.3 ($C_1$), 147.3, 150.6 ($C_{2'}$, $C_{6'}$), 169.5, 169.8 (CONH, CON). HRMS (ESI, m/z): calculated for $C_{23}H_{31}N_3O_2Na$ $[M+Na]^+$: 404.2308; found: 404.2302.

***N*$^2$-Octyl-*N*$^1$-phenyl-*N*$^2$-(pyridin-4-ylcarbonyl)-β-alaninamide (23).** Obtained from amine **26** (85 mg, 0.31 mmol), isonicotinic acid (53 mg, 0.43 mmol), EDC (67 mg, 0.43 mmol) and HOBt (58 mg, 0.43 mmol) in 43% yield (51 mg). Chromatography: hexane/EtOAc, 2:8.

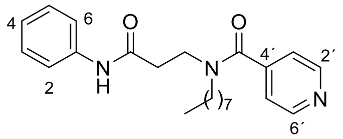

$R_f$ (EtOAc): 0.19. IR (ATR, ν): 3310 (NH), 1624 (CON), 1547 (Ar). $^1$H-NMR ($CDCl_3$, δ): amide rotamers A:B, 4:1: 0.76 (t, $J$ = 6.9 Hz, 3H, $CH_3$), 1.02-1.07 (m, 10H, $(CH_2)_5CH_3$), 1.41 (qt, $J$ = 6.9 Hz, 2H, $CH_2(CH_2)_5CH_3$), 2.40 (m, 2H, $CH_2CO$, rotamer B), 2.70 (t, $J$ = 6.5 Hz, 2H, $CH_2CO$, rotamer A), 3.12 (t, $J$ = 7.6 Hz, 2H, $(CH_2)_6CH_2N$, rotamer A), 3.41 (m, 2H, $(CH_2)_6CH_2N$, rotamer B), 3.50 (m, 2H, $COCH_2CH_2N$, rotamer B), 3.74 (t, $J$ = 6.5 Hz, 2H, $COCH_2CH_2N$, rotamer A), 7.00 (t, $J$ = 7.3 Hz, 1H, $H_4$), 7.11 (d, $J$ = 5.8 Hz, 2H, $H_{3'}$, $H_{5'}$), 7.16-7.21 (m, 2H, $H_3$, $H_5$), 7.41 (d, $J$ = 7.9 Hz, 2H, $H_2$, $H_6$), 8.55 (d, $J$ = 5.7 Hz, 2H, $H_{2'}$, $H_{6'}$), 8.78 (br s, 1H, NH). $^{13}$C-NMR ($CDCl_3$, δ): 14.0 ($CH_3$), 22.5, 26.4, 28.8, 29.0 (2C), 31.6 ($(CH_2)_6CH_3$), 35.9 ($CH_2CO$), 42.4 ($CH_2N$), 50.4 ($(CH_2)_6CH_2N$), 119.8 ($C_2$, $C_6$), 120.8 ($C_{3'}$, $C_{5'}$), 124.2 ($C_4$), 128.9 ($C_3$, $C_5$), 138.2 ($C_1$), 144.1 ($C_{4'}$), 150.3 ($C_{2'}$, $C_{6'}$), 169.2, 169.9 (CON, rotamers A and B), 171.1 (CONH). HRMS (ESI, m/z): Calculated for $C_{23}H_{31}N_3O_2Na$ $[M+Na]^+$: 404.2308. Found: 404.2311.

***N*$^2$-(1-Benzofuran-3-ylcarbonyl)-*N*$^2$-octyl-*N*$^1$-phenyl-β-alaninamide (24).** Obtained from amine **26** (100 mg, 0.36 mmol), benzofuran-3-carboxylic acid (117 mg, 0.72 mmol), EDC (138 mg, 0.72 mmol), and HOBt (97 mg, 0.72 mmol), in 57% yield (86 mg). Chromatography: DCM/MeOH, 95:5.

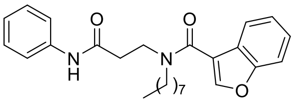

$R_f$ (DCM/ethanol 95:5): 0.38. IR (ATR, ν): 3313 (NH), 1683 (CO), 1609, 1548, 1492, 1443 (Ar). $^1$H-NMR ($CDCl_3$, δ): 0.84 (t, $J$ = 6.9, 3H, $CH_3$), 1.14-1.26 (m, 10H, $(CH_2)_5CH_3$), 1.54 (m, 2H, $CH_2(CH_2)_5CH_3$), 2.76 (m, 2H, $CH_2CO$), 3.45 (t, $J$ = 7.5, 2H, $NCH_2(CH_2)_6$), 3.85 (t, $J$ = 6.5, 2H, $NCH_2CH_2CO$), 7.06 (t, $J$ = 7.4, 1H, $H_4$), 7.15 (t, $J$ = 7.4, 1H, $CH_{benzof}$), 7.22-7.33 (m, 3H, $H_3$, $H_5$, $CH_{benzof}$), 7.46-7.61 (m, 4H, $H_2$, $H_6$, $2CH_{benzof}$), 7.74 (s, 1H, $H_3$, $H_{benzof}$), 9.15 (br s, 1H, NH). $^{13}$C-NMR ($CDCl_3$, δ): 14.0 ($CH_3$), 22.5, 26.5, 29.1 (3C), 31.7 ($(CH_2)_6CH_3$), 36.2 ($CH_2CO$), 43.0 ($NCH_2CH_2CO$), 50.4 ($NCH_2(CH_2)_6$), 111.7 ($CH_{benzof}$), 117.2 ($C_{benzof}$), 119.9 ($C_2$, $C_6$), 120.9, 123.7 ($2CH_{benzof}$), 124.1 ($C_4$), 125.3 ($CH_{benzof}$), 125.4 ($C_{benzof}$), 128.8 ($C_3$, $C_5$), 138.4 ($C_1$), 143.9 ($CH_{benzof}$), 154.6 ($C_{benzof}$), 165.4 (CON), 169.6 (CONH). HRMS (ESI, m/z): calculated for $C_{26}H_{32}N_2O_3Na$ $[M+Na]^+$: 443.2305, found: 443.2319.

***N*$^3$-(1*H*-Indol-2-ylcarbonyl)-*N*$^3$-octyl-*N*$^1$-Phenyl-β-alaninamide (25).** Obtained from amine **26** (387 mg, 1.4 mmol), 1*H*-indol-2-carboxylic acid (451 mg, 2.8 mmol), EDC (435 mg, 2.8 mmol), and HOBt (378 mg, 2.8 mmol), in 73% yield (420 mg). Chromatography: hexane/EtOAc, 7:3.

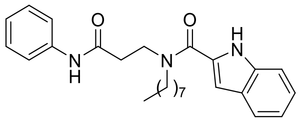

$R_f$ (hexane/EtOAc, 1:1): 0.47. mp: 142-144 ºC. IR (ATR, ν): 3284 (NH), 1666 (CO), 1600, 1533, 1442 (Ar). $^{1}$H-NMR ($CDCl_3$, δ): 0.79 (t, $J$ = 6.6, 3H, $CH_3$), 1.16 (m, 10H, $(CH_2)_5CH_3$), 1.59 (m, 2H, $CH_2(CH_2)_5CH_3$), 2.62 (t, $J$ = 6.8, 2H, $CH_2CO$), 3.50 (m, 2H, $NCH_2(CH_2)_6$), 3.80 (m, 2H, $NCH_2CH_2CO$), 6.63 (m, 1H, $CH_{indol}$), 6.92-7.03 (m, 2H, $H_4$, $CH_{indol}$), 7.09-7.16 (m, 3H, $H_3$, $H_5$, $CH_{indol}$), 7.26 (d, $J$ = 8.2 Hz, 1H, $CH_{indol}$), 7.40 (d, $J$ = 7.9 Hz, 2H, $H_2$, $H_6$), 7.51 (d, $J$ = 7.8 Hz, 1H, $CH_{indol}$), 8.53 (br s, 1H, NH), 9.82 (br s, 1H, NH). $^{13}$C-NMR ($CDCl_3$, δ): 14.1 ($CH_3$), 22.7, 26.8, 28.8, 29.2, 29.3, 31.8 ($(CH_2)_6CH_3$), 36.3 ($CH_2CO$), 44.5 ($NCH_2CH_2CO$), 49.9 ($NCH_2(CH_2)_6$), 105.2, 111.8 ($2CH_{indol}$), 120.0 ($C_2$, $C_6$), 120.6, 122.1 ($2CH_{indol}$), 124.3 ($C_4$), 124.6 ($CH_{indol}$), 127.9 ($C_{indol}$), 128.9 ($C_3$, $C_5$), 129.2, 135.7 ($2C_{indol}$), 138.1 ($C_1$), 163.4 (CON), 169.5 (CONH). MS (ESI): 420.4 [$(M+H)^+$]. Elemental analysis calculated for $C_{26}H_{33}N_3O_2$: C, 74.43; H, 7.93; N, 10.02; found: C, 74.19; H, 7.83; N, 10.00.

***N*$^2$-Octyl-*N*$^1$-phenyl-*N*$^2$-(*N*-Boc-pyrrolidin-2-ylcarbonyl)-β-alaninamide (27).** Obtained from amine **26** (71 mg, 0.26 mmol), *N*-(tert-butoxycarbonyl)-proline (110 mg, 0.51 mmol), EDC (98 mg, 0.51 mmol), and HOBt (69 mg, 0.51 mmol), in 64% yield (79 mg). Chromatography: hexane/EtOAc, 1:1.

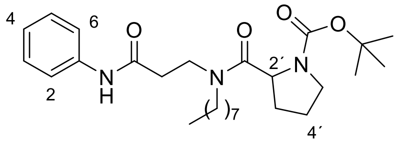

$R_f$ (hexane/EtOAc, 1:1): 0.27. IR (ATR, ν): 3312 (NH), 1685, 1667 (CO), 1604, 1546, 1441 (Ar). $^{1}$H-NMR ($CDCl_3$, δ): amide rotamers A:B, 2:1: 0.80-0.88 (m, 3H, $CH_3CH_2$), 1.17-1.28 (m, 10H, $(CH_2)_5CH_3$), 1.44-1.67 (m, 11H, $CH_2(CH_2)_5CH_3$, $C(CH_3)_3$), 1.74-1.95 (m, 2H, $CH_2CH_2CH$), 2.06-2.19 (m, 2H, $CH_2CH$), 2.31-2.51 (m, 2H, $CH_2CO$), 2.60-2.79 (m, 2H, $CH_2NBoc$, rotamer A), 3.00-3.32 (m, 2H, $NCH_2(CH_2)_6$), 3.52-3.65 (m, 2H, $NCH_2CH_2CO$), 3.67-3.87 (m, 2H, $CH_2NBoc$, rotamer B), 4.41-4.57 (m, 1H, CH), 7.01 (t, $J$ = 7.3 Hz, 1H, $H_4$), 7.26 (d, $J$ = 7.9 Hz, 2H, $H_3$, $H_5$), 7.55 (d, $J$ = 7.7 Hz, 2H, $H_2$, $H_6$, rotamer A), 7.74 (d, $J$ = 7.7 Hz, 2H, $H_2$, $H_6$, rotamer B), 9.13 (br s, 1H, NH, rotamer A), 9.34 (br s, 1H, NH, rotamer B). $^{13}$C-NMR ($CDCl_3$, δ): 14.0 ($CH_3CH_2$), 22.6 (2C), 24.4, 24.7, 26.7, 27.7 ($4CH_2$, rotamers A and B), 28.5, 28.6 ($C(CH_3)_3$, rotamers A and B), 29.1, 29.3, 30.0, 30.1, 30.2, 31.7 (2C) ($4CH_2$, rotamers A and B), 37.1, 37.9 ($CH_2CO$, rotamers A and B), 44.9, 46.0, 47.2, 47.3, 47.9, 50.0 ($3CH_2N$, rotamers A and B), 56.1, 56.5 (CH, rotamers A and B), 80.0, 80.1 ($C(CH_3)_3$, rotamers A and B), 120.1, 120.2 ($C_2$, $C_6$, rotamers A and B), 123.8, 124.0 ($C_4$, rotamers A and B), 128.7 ($C_3$, $C_5$), 138.5, 139.0 ($C_1$, rotamers A and B), 154.7, 154.9 (NCOO, rotamers A and B), 169.6, 171.2, 172.5, 173.1 (2CO, rotamers A and B). MS (ESI): 474.3 $[M+H]^+$, 374.0 $[M-Boc+H]^+$.

***N*$^2$-Octyl-*N*$^1$-phenyl-*N*$^2$-(*N*-Boc-*R*-pyrrolidin-2-ylcarbonyl)-β-alaninamide (*R*-27).** Obtained from amine **26** (100 mg, 0.36 mmol), *N*-(tert-butoxycarbonyl)-D-proline (156 mg, 0.72 mmol), EDC (139 mg, 0.72 mmol), and HOBt (98 mg, 0.72 mmol), was obtained in 74% yield (105 mg). Spectroscopic data were in agreement with those described for its racemic counterpart. Chiral HPLC-MS, $r_t$ (min): 6.66 (Chiral HPLC method A). ee> 99%

***N*$^2$-Octyl-*N*$^1$-phenyl-*N*$^2$-(*N*-Boc-*S*-pyrrolidin-2-ylcarbonyl)-β-alaninamide (*S*-27).** Obtained from amine **26** (72 mg, 0.26 mmol), *N*-(tert-butoxycarbonyl)-L-proline (109 mg, 0.50 mmol), EDC (97 mg, 0.50 mmol), and HOBt (67 mg, 0.50 mmol), was obtained in 78% yield (97 mg). Spectroscopic data were in agreement with those described for its racemic counterpart. Chiral HPLC-MS, $r_t$ (min): 7.45 (Chiral HPLC method A). ee> 99%

• ***N*$^2$-Octyl-*N*$^1$-phenyl-*N*$^2$-(pyrrolidin-2-ylcarbonyl)-β-alaninamide (14).** To a solution of *N*-Boc pyrrolidine **27** (63 mg, 0.13 mmol) in anhydrous DCM (0.5 mL), TFA (200 μL, 2.7 mmol) was added. The reaction mixture was stirred at rt for 1 h, basified with $NaHCO_3$ until pH 8 and extracted with DCM. The organic extract was washed with a saturated solution of NaCl, dried over $Na_2SO_4$, filtered, and the solvent was removed under reduced pressure, obtaining the desired final compound **14** in 70% yield (34 mg).

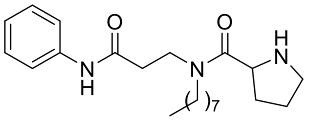

$R_f$ (EtOAc/methanol, 7/3): 0.19. IR (ATR, ν): 3269 (NH), 1644 (CO), 1605, 1548, 1494, 1444 (Ar). $^1$H-NMR ($CDCl_3$, δ): amide rotamers A:B, 2:1: 0.83-0.88 (m, 3H, $CH_3$), 1.21-1.24 (m, 10H, $(CH_2)_5CH_3$), 1.45-2.30 (m, 6H, $CH_2(CH_2)_5CH_3$, $(CH_2)_2CH$), 2.56-3.05 (m, 4H, $CH_2CO$, $CH_2NH$), 3.15-3.34 (m, 2H, $NCH_2(CH_2)_6$), 3.55-3.76 (m, 2H, $NCH_2CH_2CO$), 3.91 (t, *J* = 7.5 Hz, 1H, CH, rotamer A), 4.37 (t, *J* = 7.5 Hz, 1H, CH, rotamer B), 5.10 (br s, 1H, NH), 7.05 (t, *J* = 7.3 Hz, 1H, $H_4$), 7.23-7.30 (m, 2H, $H_3$, $H_5$), 7.56 (d, *J* = 7.7 Hz, 2H, $H_2$, $H_6$, rotamer A), 7.61 (d, *J* = 7.7 Hz, 2H, $H_2$, $H_6$, rotamer B), 8.97 (br s, 1H, NH, rotamer A), 9.58 (br s, 1H, NH, rotamer B). $^{13}$C-NMR ($CDCl_3$, δ): 14.1 ($CH_3$), 22.6, 25.8, 26.3, 26.8, 27.6, 29.1, 29.2 (2C), 29.3 (2C), 29.7, 30.6, 31.1, 31.7 ($(CH_2)_6CH_3$, $(CH_2)_2CH$, rotamers A and B), 36.3, 37.2 ($CH_2CO$, rotamers A and B), 43.2, 44.0 ($NCH_2CH_2CO$, rotamers A and B), 46.8, 46.9, 47.4, 48.3 ($NCH_2(CH_2)_6$, $CH_2NH$, rotamers A and B), 57.8, 58.2 (CH, rotamers A and B), 119.9 ($C_2$, $C_6$), 124.1 ($C_4$), 128.8, 128.9 ($C_3$, $C_5$), 138.4 ($C_1$), 168.7, 169.5, 171.4, 173.9 (2CO, rotamers A and B). HRMS (ESI, m/z): Calculated for $C_{22}H_{36}N_3O_2$ $[M+H]^+$: 374.2796; found: 374.2802.

***N*$^2$-Octyl-*N*$^1$-phenyl-*N*$^2$-(*R*-pyrrolidin-2-ylcarbonyl)-β-alaninamide (*R*-14).** Obtained from (*R*)-**27** (75 mg, 0.16 mmol) and TFA (244 µL, 3.2 mmol) in 90% yield (59 mg). Spectroscopic data were in agreement with those described for racemic compound **14**. Chiral HPLC-MS, $r_t$ (min): 5.99 (Chiral HPLC method B). ee> 99%. $[\alpha]^D_{20}$: -103.2° (c = 0.4, methanol).

***N*$^2$-Octyl-*N*$^1$-phenyl-*N*$^2$-(*S*-pyrrolidin-2-ylcarbonyl)-β-alaninamide (*S*-14).** Obtained from (*S*)-**27** (90 mg, 0.19 mmol) and TFA (293 µL, 3.8 mmol) in 70% yield (50 mg). Spectroscopic data were in agreement with those described for racemic compound **14**. Chiral HPLC-MS, $r_t$ (min): 10.18 (Chiral HPLC method B). ee> 99%. $[\alpha]^D_{20}$: + 100.8° (c = 0.4, methanol).
